# DNA Methylation profiles of diverse *Brachypodium distachyon* aligns with underlying genetic diversity

**DOI:** 10.1101/039602

**Authors:** SR Eichten, T Stuart, A Srivastava, R Lister, JO Borevitz

## Abstract

DNA methylation, a common modification of genomic DNA, is known to influence the expression of transposable elements as well as some genes. Although commonly viewed as an epigenetic mark, evidence has shown that underlying genetic variation, such as transposable element polymorphisms, often associate with differential DNA methylation states. To investigate the role of DNA methylation variation, transposable element polymorphism, and genomic diversity, whole genome bisulfite sequencing was performed on genetically diverse lines of the model cereal *Brachypodium distachyon*. Although DNA methylation profiles are broadly similar, thousands of differentially methylated regions are observed between lines. An analysis of novel transposable element indel variation highlighted hundreds of new polymorphisms not seen in the reference sequence. DNA methylation and transposable element variation is correlated with the genome-wide amount of genetic variation present between samples. However, there was minimal evidence that novel transposon insertion or deletions are associated with nearby differential methylation. This study highlights the importance of genetic variation when assessing DNA methylation variation between samples and provides a valuable map of DNA methylation across diverse re-sequenced accessions of this model cereal species.

**Reviewer Link to deposited data:** All data is publicly available in the NCBI short read archive under BioProject PRJNA281014. Data tables are available for download at https://drive.google.com/file/d/0BzBxfoxlBCneNkY1TFJDU29iSUU/view?usp=sharing

## Introduction

Individuals of a species are often classified based on genetic variation found between them. In addition to genetic variation, interest has grown as to other possible sources of heritable variation between individuals. Of these, methylation of cytosine residues (DNA methylation) act as an epigenomic mark that largely targets transposable elements and other repetitive sequence of the genome to prevent transposition and possibly silence cryptic promoters (Law and Jacobsen 2010; Weigel & Colot, 2012; Diez et al., 2014; Kim & Zilberman, 2014; Matzke & Mosher 2014). The function of DNA methylation also appears to impact gene expression largely through downregulation via promoter methylation (Bucher et al., 2012), or possible upregulation via gene body methylation (Zilberman & Henikoff, 2007; Teixeira & Colot, 2009; Maunakea et al., 2010). DNA methylation is one of a number of genome modifications that may be able to create an ‘epiallele’ that can be inherited independently of any underlying genetic variation (Eichten et al., 2014). Although a number of genome scale analyses of DNA methylation and its relationship to genome variation, chromatin modifications, and transcription have been undertaken in plants (Zilberman et al., 2007; Lister et al., 2008; Cokus et al., 2008; Schmitz et al., 2011; Miura et al., 2012; Chodavarapu et al., 2012; Zhong et al 2013; Eichten et al., 2013; Stroud et al., 2013; Regulski et al, 2013; Schmitz et al., 2013), the relationship of DNA methylation relative to other classes of genetic variation and patterns of genomic organization within and among species, is still emerging (Seymour et al. 2014, Dubin et al., 2015).

Advances in DNA methylation profiling have allowed a number of plant species to be profiled at the whole-genome level (Zilberman et al., 2007; Lister et al., 2008; Cokus et al., 2008; Schmitz et al., 2011; Miura et al., 2012; Chodavarapu et al., 2012; Zhong et al 2013; Eichten et al., 2013; Stroud et al., 2013; Regulski et al, 2013; Schmitz et al., 2013) leading to a basic understanding of the broad patterns of DNA methylation within the genome. The model cereal, *Brachypodium distachyon* provides a unique plant system to study DNA methylation. With a small diploid genome (∼271Mb), high genetic diversity (Vogel et al., 2009), global distribution (Garvin et al., 2008), and close relation to barley and wheat (Draper et al., 2001), it provides a unique and important model system to investigate the function of DNA methylation. Brachypodium is a monocot with a genome size that is highly amenable to sequencing analyses compared to crop systems such as maize (Schnable et al., 2009), barley (Mayer et al, 2012), or wheat (Mayer et al, 2014). To complement recent genomic sequencing efforts in this model species (IBI, 2010; Gordon et al., 2014), an understanding of the Brachypodium chromatin landscape can provide insights as to the potential effects of transposable element insertions, chromatin accessibility, and functional consequences of differential methylation in this globally diverse plant system.

DNA methylation in plants shows strong regional placement to target transposable element sequences for repression while also targeting other non-repeat sequences within the genome (Zilberman et al., 2007; Lister et al., 2008; Cokus et al., 2008; Schmitz et al., 2011; Miura et al., 2012; Chodavarapu et al., 2012; Zhong et al 2013; Eichten et al., 2013; Stroud et al., 2013; Regulski et al, 2013; Schmitz et al., 2013). Gene body methylation has been compared between orthologous genes in plants indicating a strong conservation of this intragenic methylation (Takuno & Gaut, 2013) across species. Recent evidence in Brassicaceae has also shown that DNA methylation variation between species is tied to regions of genomic variability driven largely by transposable elements (Seymour et al., 2014). As a major function of DNA methylation in plants is as a repressor of transposable element activity (Kim & Zilberman, 2014), DNA methylation is highly correlated to the positions of known transposable elements within genomes. Beyond targeting repetitive elements directly, DNA methylation can often ‘spread’ outside of the element boundary and influence nearby sequence (Hollister & Gaut, 2009; Hollister et al., 2011; Ahmed et al., 2011; Eichten et al., 2012). From this it is believed that differentially methylated regions (DMRs) can be driven by transposon insertion polymorphisms in which the presence or absence of an element dictates methylation levels in flanking low-copy sequence. Transposon polymorphism is a likely driver of genetically-controlled DNA methylation variation, however the ability to identify novel TE insertions, and therefore prospective methylation variants, has been limited to date (van Opijnen & Camilli, 2013; Nakagome et al., 2014).

To investigate the DNA methylation landscape of *Brachypodium distachyon*, we profiled seven recently resequenced lines (Gordon et al., 2014) by Whole Genome Bisulfite Sequencing (WGBS, by post bisulfite adapter tagging,PBAT) to obtain base-pair resolution profiles of DNA methylation throughout their genomes. Differentially methylated regions (DMRs) across the different DNA methylation sequence contexts (CG, CHG, and CHH, where H represents A, C, or T) are prominent across these lines when contrasted to the Bd21 reference genome. Genetic diversity in the form of SNPs and newly-identified transposable element insertion polymorphisms between lines, aligns with many of the identified DMRs, highlighting the interaction between genetic diversity and chromatin states within the species.

## Results

### DNA methylation patterns of the Bd21 reference genome

To investigate DNA methylation patterns in the *Brachypodium distachyon* reference genome Bd21, WGBS via PBAT was performed and 100 bp reads were aligned resulting in ∼22 million unique alignments with an average coverage of 8.1× (Sup Table 1). Although the sequencing data provides base-pair resolution of cytosine methylation, broader patterns of DNA methylation encompassing multiple neighboring cytosines can highlight the importance of DNA methylation in maintaining chromatin states throughout the genome. Therefore, average DNA methylation patterns across the chromosomes using non-overlapping 100 bp tiles were calculated for each of the three sequence contexts. The Bd21 reference genome contains ∼2.4-2.7 million tiles with at least a single cytosine for each respective methylation context (Figure 1A). The WGBS performed provides coverage across ∼90% of all possible tiles for each methylation context. DNA methylation levels across genomic tiles display a largely bimodal distribution for CG and CHG methylation levels (Supplemental Figure 1). From this, tiles were called either ‘methylated’ or ‘unmethylated’ based on these distributions, requiring >50% weighted methylation (Schultz et al., 2012) for CG, 30% for CHG, and 10% for CHH. The proportion of methylated tiles for CHH methylation is much smaller than the CG and CHG contexts (Figure 1A). When viewing methylation levels across chromosomes, higher levels of CG and CHG methylation are visible in the gene sparse and repeat rich pericentromeric regions (Figure 1B). This is in contrast to CHH methylation which, although an order of magnitude less abundant than CG or CHG methylation, displays a more uniform distribution across the chromosome (Figure 1B).

**Figure 1:**
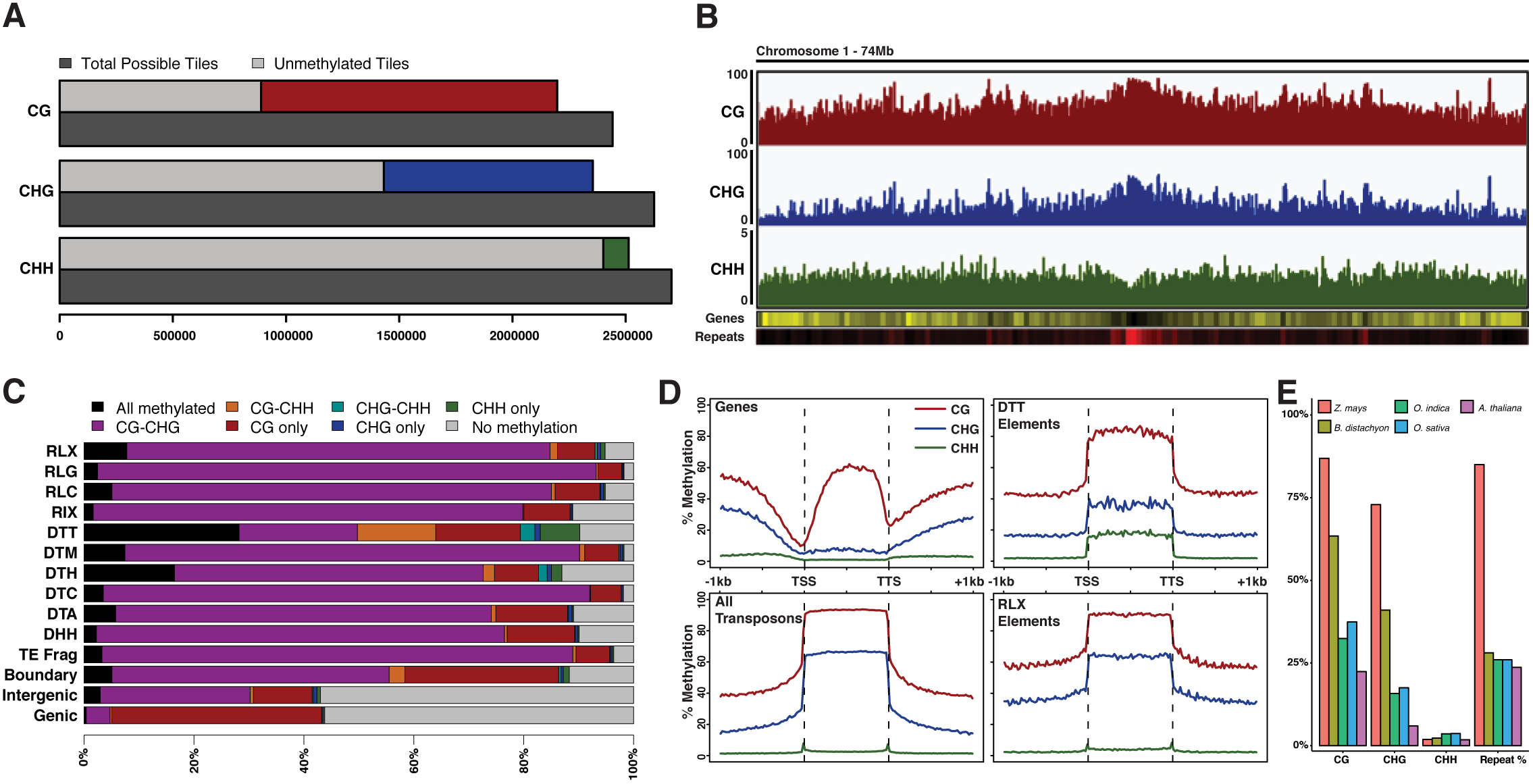
Genomic profiles of DNA methylation of Brachypodium distachyon. (A) Total number of 100 bp genomic tiles (dark grey) are compared to the number of genomic tiles with sequence coverage for all three methylation sequence contexts. Coverage bar is split between unmethylated tiles (light grey) and methylated tiles (red = CG; blue= CHG; green=CHH). (B) Chromosome profile of average methylation in CG, CHG, and CHH contexts. Gene density (yellow high) and repeat density (red high) are also shown. (C) Percentage barplot of genomic tile annotation states within the genome. Bars are divided by eight possible Bd21 methylation states. (D) Relative methylation of annotated genes, all transposons, DTT transposons only, and RLX transposons only. Flanking 1 kb of genes and repetitive elements is also shown outside dashed vertical lines. (E) Average level of CG, CHG, CHH, and proportion of genomes consisting of repetitive sequences across the genome of various grass species.

With each genomic tile classified based on its methylation type, a total of eight possible methylation states, ranging from no methylation to fully methylated in all contexts, can be identified for windows containing coverage across all cytosine contexts. The majority of genomic tiles display no methylation, with the remaining tiles largely falling into ‘CG-only’ and ‘CG-CHG’ methylation states (Sup Fig 2). DNA methylation patterns may differ across different genomic features such as genes and transposable elements. Therefore, each genomic tile was grouped based on its nearest intersecting annotated feature (Sup Fig 3). 28% of genomic tiles are found in TEs, 33% in genes, 37% are intergenic, and 2% overlap both genes and TEs. The type of DNA methylation within genes and transposon families provides evidence for different methylation profiles for different features (Figure 1C). Genic and intergenic genomic tiles are frequently unmethylated compared to transposable element regions. Genic methylation is largely CG-only with intergenic regions displaying CG-only and CG-CHG tiles predominantly. Transposable elements are largely methylated, with CG-CHG only tiles being most common. These patterns are consistent with other studies in plants, however the DTT class of Sub1 / Mariner elements shows a unique pattern of methylation with an increase in CHH methylation compared to other TEs (Figure 1C; Supplemental Figure 4). Genomic tiles that intersect the boundaries of both genes and TEs contain a combination of methylation states found in both genes and TEs. Overall, DNA methylation patterns are substantially different between annotation features, with DTT Mariner elements constituting a proportionally large source of CHH methylation within the genome (∼8% of all CHH-containing tiles in the genome are found in the ∼1% of genomic tiles that are DTT elements).

DNA methylation is known to target genes and transposable elements differently. Previous reports of DNA methylation within *Brachypodium distachyon* have indicated a high proportion of CG methylation within annotated gene bodies (Takuno & Gaut, 2013). When comparing all annotated genes in the Bd21 reference genome, a clear pattern of CG gene body methylation was apparent with minimal levels of CHG and CHH methylation (Figure 1D). This body methylation was maintained with intronic sequence removed, indicating gene body methylation is not a byproduct of intron content (Sup Fig 5). There is a sharp decrease in CG and CHG methylation surrounding the transcription start site. In contrast, methylation patterns are strikingly different when averaged across genomic repeats in the Bd21 reference genome (Figure 1D). CG methylation is close to a saturating level across repetitive elements, and CHG methylation is also increased compared to genes, though to a lesser extent. For both genes and repeats, DNA methylation appears to return to genomic-average levels once outside the boundaries of the annotated features (Figure 1D). When genes and transposable elements were divided based on size, CG gene body methylation shows distinct increases positively correlated with gene size leading to minimal methylation for genes smaller than 2 kb (Sup Fig 6). In contrast, repetitive elements of all sizes display high levels of CG methylation, with increased levels in smaller elements with CHG and CHH showing decreasing levels as repeat size increases (Sup Fig 6). DNA methylation levels over DTT elements display elevated levels of CHH as indicated via tile analysis (Figure 1D). This is in contrast to other repeat superfamilies such as RLX which do not show elevated levels within the element boundaries.

Plant genomes differ greatly in their repetitive DNA content (i.e. transposable elements). A comparison of *Brachypodium distachyon* global methylation levels to other plant species was performed using published WGBS reads, analyzed with the same alignment parameters (see methods). When comparing average methylation level to the published repeat content of each genome (Schnable et al., 2009; IBI, 2010; Yu et al., 2002; Goff et al., 2002; Maumas & Quesneville, 2014), there is a clear positive association between repeat content, CG, and CHG methylation (Figure 1E). In contrast, asymmetric CHH methylation does not appear to correlate with genomic repeat proportion. Bd21 displays 2.3% genomic CHH methylation compared to 1.8% in *Arabidopsis thaliana* and 3.7% in *Oryza sativa*. Curiously, Bd21 displays a higher proportion of CG and CHG methylation relative to its genomic repeat content compared to other species. Genic DNA methylation appears similar between species (Sup Fig 7) with similar patterns of CG gene body methylation for all contexts.

Initial genome sequencing of the Bd21 reference line showed that chromosome 5 displayed the lowest gene density, and increased retrotransposon density, compared to other chromosomes (IBI 2010). Given the positive relationship between repeat content and methylation level, overall DNA methylation levels were compared between chromosomes (Sup Fig 8). As expected, chromosome 5, with the highest repetitive content, displayed the highest levels of CG and CHG methylation compared to other chromosomes. Overall, DNA methylation in Bd21 follows similar patterns of methylation that have been observed in other plant species (Lister et al., 2008; Cokus et al., 2008; Chodavarapu et al., 2012; Eichten et al., 2013; Mirouze & Vitte, 2014), with a slight increase in overall symmetric methylation levels given its genomic repeat content.

### Differential methylation across diverse B. distachyon samples

DNA methylation is known to vary between individuals of a species (Vaughn et al., 2007; Zhang et al., 2008; Eichten et al., 2013; Schmitz et al., 2013) and can act as a source of heritable variation impacting gene expression (Stokes et al., 2002; Zilberman et al., 2007; Vaughn et al., 2007; Becker et al., 2011; Schmitz et al., 2011). To investigate DNA methylation variation within *Brachypodium distachyon*, six additional WGBS profiles were created for the inbred lines Bd21-3, Bd3-1, Bd30-1, BdTR12c, Koz-3, and Bd1-1 (Supplementary Table 1). These lines were chosen as they have recently been sequenced as additional reference lines and are commonly used divergent strains within the species (Gordon et al., 2014; Vogel et al., 2009). Reads obtained for each sample were aligned to the SNP corrected reference genomes corresponding to the selected inbred lines (Gordon et al., 2014). Overall patterns of methylation are highly similar between lines (Sup Fig 9) indicating limited broad-scale variation. The resulting methylation data was used to identify differentially methylated regions (DMRs) between these lines.

The impact of qualitative differential methylation within the genome appears to have more functional consequences when viewed as regions of multiple cytosines rather than single sites. Therefore, DNA methylation for all three contexts was averaged independently across non-overlapping 100 bp windows. Also, as methylation levels between sequence contexts are maintained through different mechanisms (Law & Jacobsen, 2010) it is important to classify differential methylation for each context independently. A series of filters were used to classify a DMR window as a CG, CHG, or CHH DMR: for each genotype pairwise comparison, differential methylation was calculated as two or more concurrent windows with at least 3x coverage between both lines and at least two cytosines with coverage that have a difference in methylation of at least 70% (CG) or 50% (CHG; see methods). After collapsing adjacent DMR windows, 5,588-9,550 CG-DMRs and 4,568-8,396 CHG-DMRs were identified across the genome for each line when compared to the Bd21 reference (Figure 2A; Supplementary Table 2). CHH methylation is present at much lower abundances than CG/CHG (Figure 1A-B) and therefore requires separate criteria for DMR discovery. CHH DMRs were classified as two or more concurrent windows with at least 3x coverage and at least 8 cytosines with coverage within the analyzed windows. Differential methylation required one sample to display ‘low’ (<= 5%) and the other ‘high’ (>=20%) methylation to be considered a DMR. In total, 520-921 CHH DMRs were identified between the diverse lines and Bd21 (Figure 2G; Supplementary Table 3).

**Figure 2:**
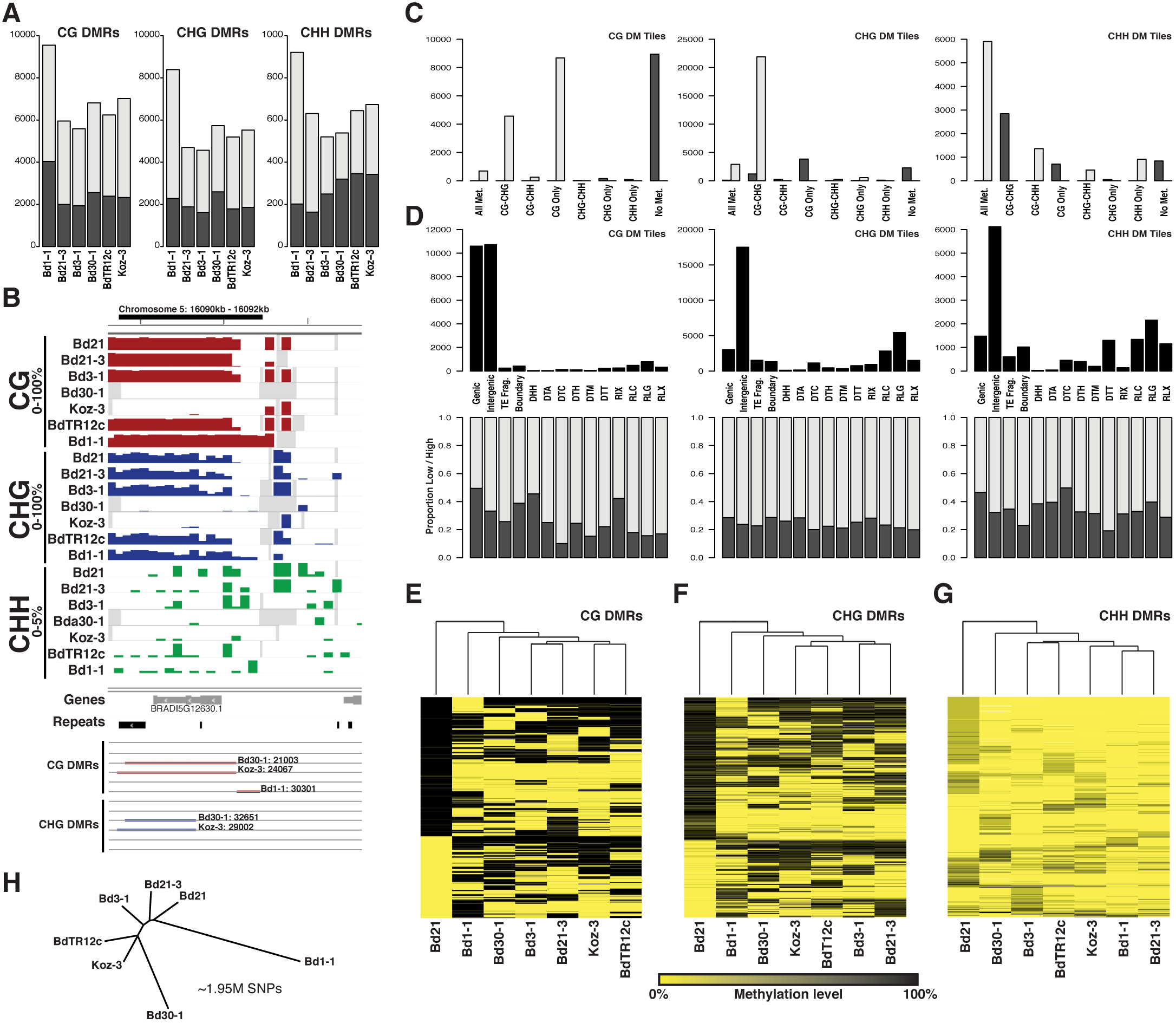
Differential methylation across Brachypodium inbred lines. (A) Total number of CG, CHG, and CHH DMRs classified for each of the six diverse inbreds. Grey and black indicate low and high methylation compared to Bd21 respectively. (B) An example DMR region with 100 bp averaged methylation profiles for all seven inbreds. Gene and repeat models are provided below. Bd30-1, Koz-3, and Bd1-1 CG/CHG DMRs are highlighted. (C) Number of differentially methylated tiles across eight Bd21 methylation classes. Grey and black indicate low and high methylation compared to Bd21 respectively. (D) The total number of DM tiles (top) and proportion of low (grey) and high (black) methylation for DM tiles (bottom) across genomic annotation classes. (E-G) Hierarchical clustering of all identified CG, CHG, and CHH DMRs. Dendrogram indicates overall similarity between samples. Given all DMRs discovered against Bd21, Bd21 acts as an outgroup in this analysis. (H) Neighbor-Joining tree of the seven reference lines based on 1.95 million SNPs with calls for all samples. SNPs encoded as Bd21 or alternate.

For all three methylation sequence contexts, the vast majority (∼99%) of genomic tiles do not display differential methylation. However, differential methylation was apparent in certain regions of the genome (Figure 2B). We observed that differentially methylated (DM) tiles often display changes only in specific DNA methylation state combinations (Figure 2C; Bd1-1 shown as example). For CG DM tiles, high methylation (when compared to the Bd21 reference) was almost exclusively found in tiles without methylation in any context for Bd21. Low CG methylation compared to Bd21 is most common across Bd21 tiles displaying CG, or both CG and CHG (CG-CHG), methylation. CHG DM tiles appear to come from a largely different set of genomic tiles with high CHG methylation events occurring most often in Bd21 regions displaying CG methylation. Low CHG methylation occurs for tiles where Bd21 displays CG-CHG methylation. High CHH methylation DM tiles also are most common where Bd21 displays CG-CHG methylated tiles. Surprisingly, low CHH methylation is predominantly derived from genomic tiles where Bd21 is methylated in all three sequence contexts (Figure 2C). Overall differential methylation across the three sequence contexts appears to be derived from different chromatin states when considering their methylation state in Bd21.

Beyond comparisons of methylation state, DM tiles are found in different genomic features depending on sequence context (Fig 2C; Bd1-1 shown). CG DM tiles are almost all found in genic or intergenic regions. Very few DM tiles are found in any transposable element classes. However, low CG DMR tiles appear enriched for most transposable element classes compared to the overall number of low DM tiles (60%; Figure 2D bottom graph). CHG DM tiles most commonly display a low methylation state compared to the reference (75%). CHG DM tiles are found within intergenic regions most often with no evidence of annotation-specific enrichment of DM direction. CHH DM tiles, although predominantly intergenic, are much more common across transposable elements. The majority of annotated features display a similar proportion of methylation differences for CHH DM tiles. Although the Bd1-1 DM tiles are shown (Figure 2C,D), these patterns appear conserved across the other five lines compared (data not shown).

Individual 100 bp tiles can further be collapsed into larger regions of differential methylation (see methods). DMRs were found throughout all five chromosomes and were slightly depleted within gene-poor regions of the genome (Sup Fig 10). DMRs ranged in size from 200 bp (minimum size allowed) to as large as 4.7 kb. CHH DMRs tend to be much shorter than CHG or CG DMRs (Sup Fig 11). DMRs between lines displayed considerable overlap, with over 30% of CG-DMRs found in one line at least partially overlapping with a DMR from a different comparison. Across the three context-specific types of DMRs identified, patterns highlighting the relationship between sequence contexts were easily identified (Sup Fig 12). The majority of CG DMRs (65%) also display differential methylation in the CHG context (Sup Fig 12). CHG DMRs show a similar pattern in respect to CG methylation, however only ∼23% of CHG DMRs also display differences in CG methylation. Of the CHG DMRs that do not display differences in CG methylation state, almost all (∼99%) have high levels of CG methylation. CHH DMRs showed a less distinct pattern of increased methylation in other contexts. CG and CHG methylation is largely stable in CHH DMR regions (Sup Fig 12).

DNA methylation states for each DMR were calculated for all seven inbreds to identify common states that are shared between lines (Figure 2B; Supplementary Table 2). To investigate the relationship between samples, hierarchical clustering of DMR states across all samples was performed (Figure 2E-G). Clustering indicates that many DMRs display differential methylation in only one or two of the seven samples, and may be rare variants unique to the individual line. Beyond Bd21, which acts as a pseudo out-group given its relationship to DMR identification, Bd1-1 is the most diverged line compared to the rest for both CG and CHG DMR sets. This, combined with Bd1-1 displaying the highest number of DMRs for all contexts (Figure 2A) indicates that Bd1-1 contains the most diverged chromatin state compared to the other examined lines. This is consistent with the genetic relationship between lines as seen when constructing a neighbor-joining tree from over two million filtered SNPs (Figure 2H; Gordon et al., 2014).

### Biological replicates quantify heritable methylation variation between genotypes

Many attempts to profile absolute DNA methylation levels have often relied on single replicate data and qualitative analysis, as presented above, due to high sample cost. However, single-replicate data does not allow for the direct measurement of methylation variability between biological replicates and therefore limits the ability to robustly identify DMRs that consider intra-genotype variation. Recent work investigating Bd21 methylation patterns has shown that there is a level of biological variability between biological replicates of the same inbred line (Roessler et al., 2015), highlighting the need to acknowledge intra-genotype variability and stability of DNA methylation.

To investigate the biological variability of DNA methylation within an inbred line, 14 additional samples consisting of five replicates of Bd21, Bd1-1, and four replicates of Bd3-1, were analyzed by WGBS (Supplemental Table 4; see methods). Overall methylation values correlated highly (r2 ∼0.9) with single-replicate data of the previous experiment and replicate samples showed high conservation for CG and CHG methylation states (Figure 3A-B). CHH methylation appeared to be more variable with overall lower correlation coefficients (R2∼0.35) when compared to the other methylation contexts (Figure 3C).

**Figure 3:**
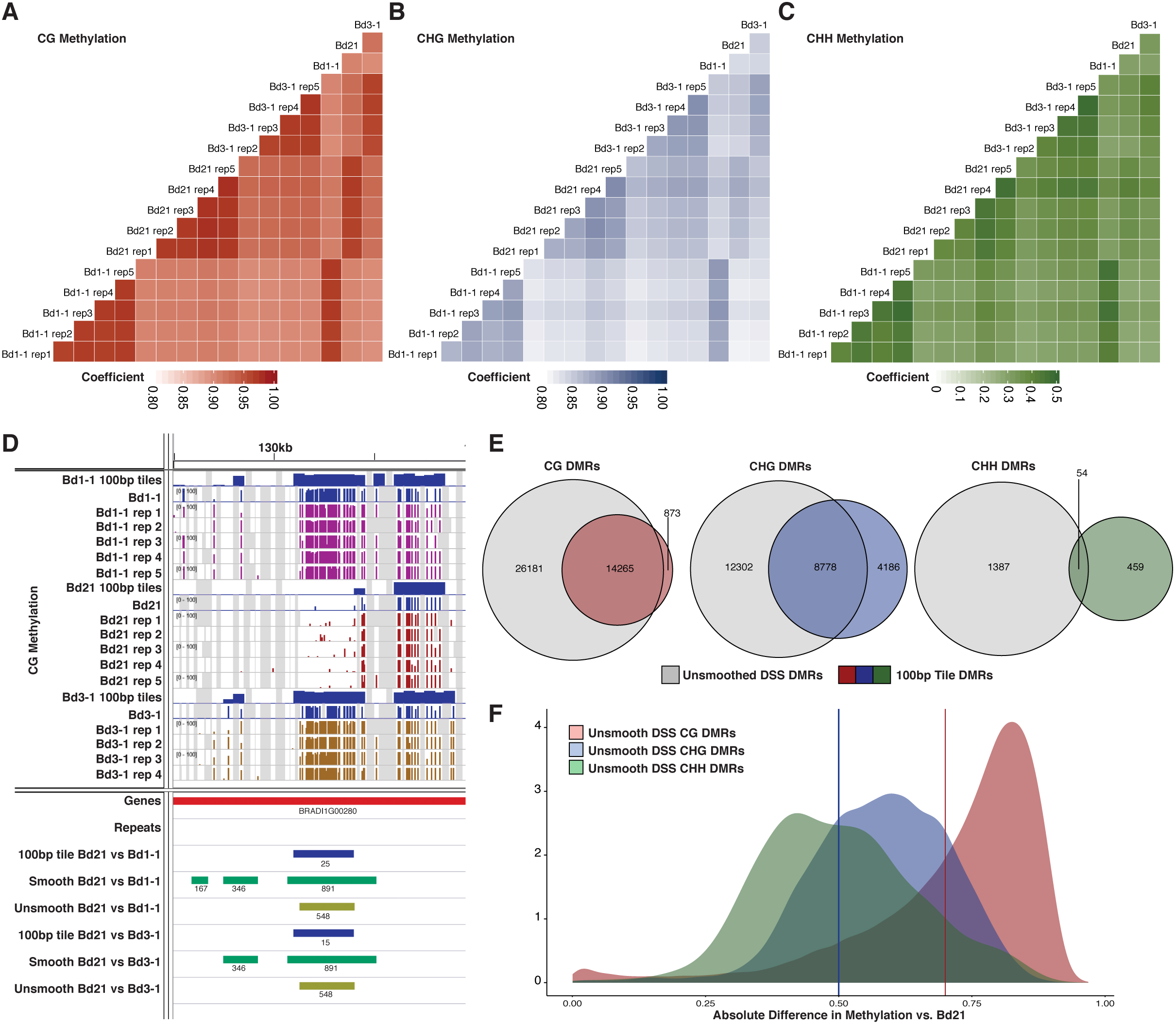
Comparisons of biological replicates of whole genome bisulfite sequencing. (A-C) Pearson correlation plots of all genomic cytosines with coverage for (A) CG, (B) CHG, and (C) CHH methylation. Note different scale for CHH methylation compared to other two contexts. (D) Genome view of a CG DMR region highlighting the 100 bp windowing (Blue), Smooth DSS (green), and unsmooth DSS (yellow) DMRs called. Tracks of cytosine methylation values are shown along with matching experimental sample and resulting 100bp window track. (E) Venn diagrams of unsmoothed DSS-based DMRs compared to 100 bp tile-base DMRs for CG, CHG, and CHH DMR sets. All DMRs for both Bd1-1 and Bd3-1 comparisons to the reference were included. (F) Density distribution of methylation differences between accession groups in unsmoothed DSS DMRs. Red vertical bar indicates fixed cutoff for 100 bp CG DMRs. Blue vertical bar indicates fixed cutoff for 100 bp CHG DMRs.

Pairwise differential methylation between the three replicated inbreds was evaluated using the Dispersion Shrinkage for Sequencing data method (DSS; Feng et al., 2014), which takes into account both the within and between line variation to develop a biologically robust measure of methylation variation at each cytosine position. The resulting data was then collapsed into DMRs under the default package parameters to identify regional DNA methylation variation (Figure 3D). The DSS method of analysis allows for the imputation of missing data by smoothing based on surrounding cytosine levels. When smoothing is allowed, an order of magnitude increase in the number of DMRs was found (Table 1). DMRs varied in size, with a median size of approximately 1,500 bp (Supplemental Table 5). Smoothed DSS data produces a large number of DMRs, but many of these appear to be called over regions completely lacking coverage in one of the replicated genotypes, and these are unlikely to be true DMRs but rather due to genetic divergence and difficulties with read mapping (Supplemental Figure 13). In order to limit the imputation of inflated DMR numbers, DSS was run without smoothing and methylation count data for all replicates was required for each inbred line examined. This resulted in roughly half the number of CG and CHG DMRs and a substantial reduction in CHH DMRs being called compared to the smoothed data (Table 1; Supplemental Table 6).These unsmoothed DMRs were used for further investigation.

**Table 1:**
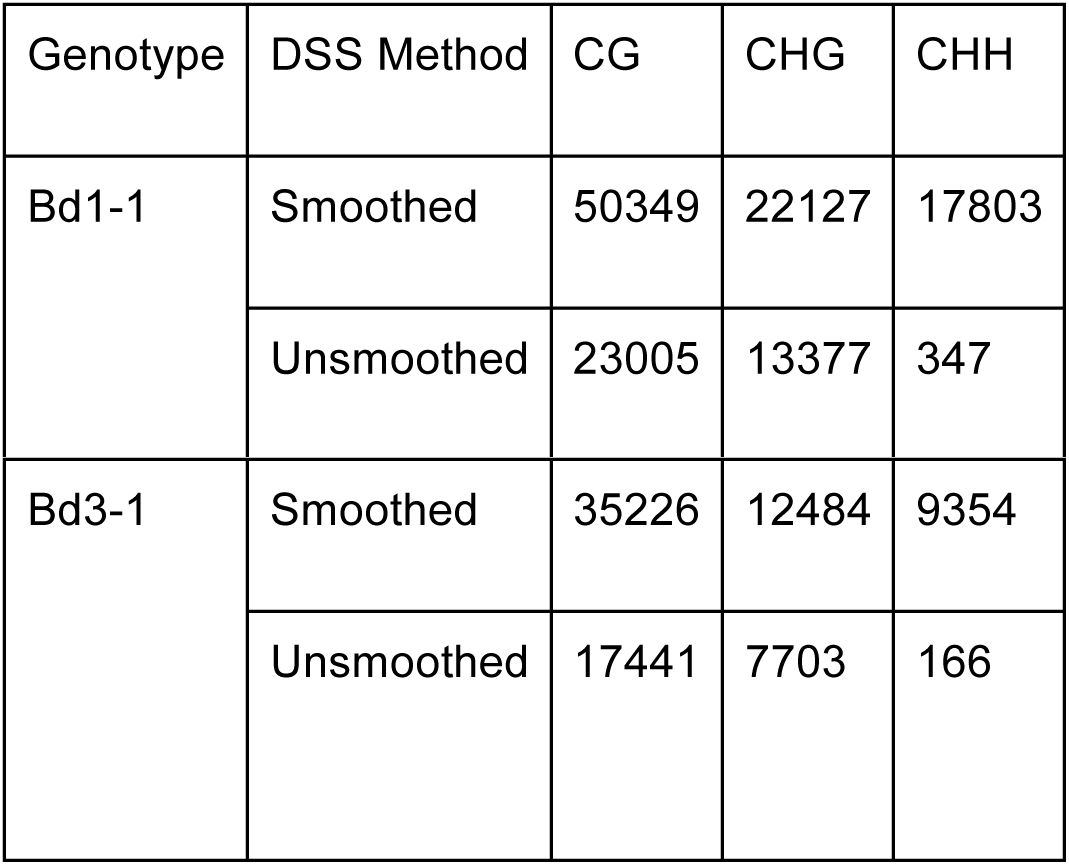
Number of DMRs called against the reference

| Genotype | DSS Method | CG | CHG | CHH |
| --- | --- | --- | --- | --- |
| Bd1-1 | Smoothed | 50349 | 22127 | 17803 |
|  | Unsmoothed | 23005 | 13377 | 347 |
| Bd3-1 | Smoothed | 35226 | 12484 | 9354 |
|  | Unsmoothed | 17441 | 7703 | 166 |

To investigate the similarities between quantitative DMRs called using DSS and qualitative DMRs from 100 bp tiles, the datasets were intersected (Figure 3E). For CG DMRs, ∼94% (14,265 of 15,138) of the combined Bd1-1 and Bd3-1 100 bp DMRs intersected with unsmoothed CG DMRs. However, ∼60% of unsmoothed CG DMRs are novel and missed from the qualitative analysis. Overlapping DMRs show a very different pattern for CHG and CHH DMRs with ∼68% and ∼4% of 100 bp DMRs intersecting the unsmooth DSS DMRs respectively. Overall, there are many DMRs unique to the unsmoothed DSS method, and minimal overlap for CHG and CHH DMRs between methods, highlighting the importance of accounting for biological variation in methylation levels when calling DMRs. This is particularly true for the more variable CHG and CHH contexts.

Although DNA methylation is a binary state at an individual cytosine, methylation levels are identified from a pool of cell types and lead to proportional differences. The DNA methylation level could be considered a quantitative trait rather than a binary state, as DSS analysis assumes. The unsmoothed DSS DMRs largely highlight differences in methylation of ∼80% for CG methylation, ∼30-80% for CHG, and ∼20-90% for CHH (Figure 3F). The majority of DMRs being called by the unsmoothed DSS method display methylation differences at, or beyond, the filtering requirements of the 100 bp CG or CHG DMRs (Figure 3F vertical bars). However there are 13,412 (∼33%) CG, 5,640 (∼27%) CHG, and 116 (∼23%) CHH DMRs from the unsmoothed DSS method that are called as significant differences which would have been omitted from the tile-based analysis as methylation differences are lower than the required threshold. These additional DMRs highlight the value of replicate data in providing additional power to identify variants with smaller overall changes in methylation.

### DNA methylation variation is correlated with genetic variation between lines

Recent evidence has shown that chromatin marks such as DNA methylation are often driven by genetic variation within the genome (Lisch and Bennetzen, 2011; Fedoroff et al., 2012; Rebollo et al., 2012; Weigel & Colot, 2012, Eichten et al., 2013). Single nucleotide polymorphism (SNP) rates (Gordon et al., 2014) were compared with 100 bp tile-based DMR frequency throughout the genome (Figure 4). DMRs for all three sequence contexts were more prevalent in genomic regions containing higher genetic diversity (Figure 4A-C; Supplemental Figure 14). These trends were present for each individual sample with stronger correlation values for CG (r2:0.314-0.699) and CHG (r2:0.279-0.541) compared to CHH (r2:0.063 - 0.545) which may be due to limited CHH DMR number (Supplemental Figure 14).

**Figure 4:**
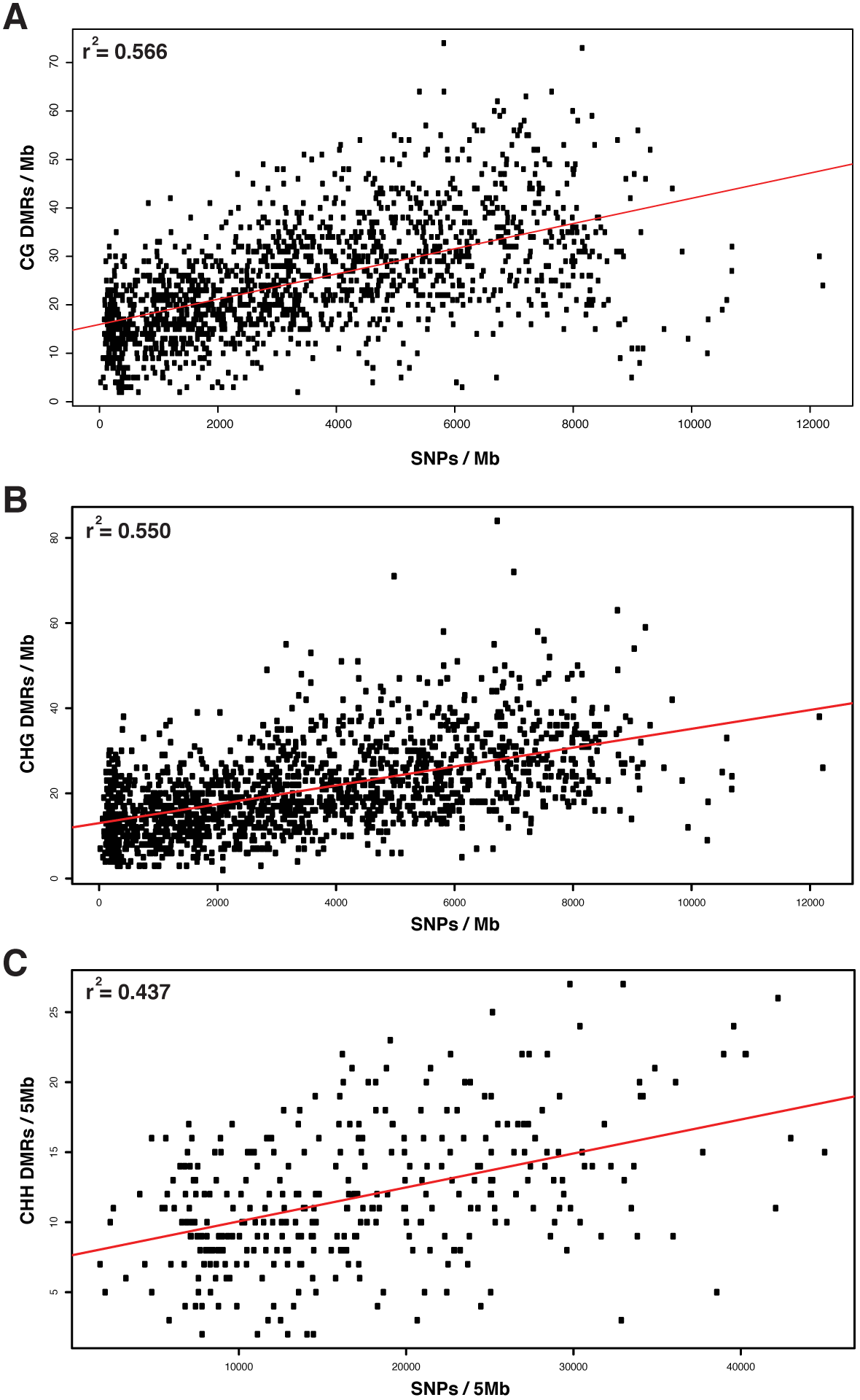
DMR density correlated with SNP density across (A) CG, (B) CHG, and (C) CHH sequence contexts. CG and CHG calculated over non-overlapping 1 Mb windows. CHH density calculated across 5Mb windows given fewer CHH DMRs.

Although there is a relationship between the local frequency of DNA methylation variation and genetic variation, this does not appear to fully explain the presence of DMRs within the genome. Across all sample contrasts, 873 DMRs are identified in low-diversity regions (>5,000 bp per SNP). The correlations observed indicate a similar relationship to previous studies in maize for which ∼50% of DMRs appear locally associated with SNP state (Eichten et al., 2013). When using unsmoothed DSS DMRs (Sup Table 6), similar patterns are observed with lower correlation values (maximum of 0.39) compared to the 100 bp tile-based DMRs (data not shown).

### Frequent transposable element polymorphisms create novel genetic diversity within the genome

Transposable elements (TEs) are often targets of DNA methylation (Lister et al., 2008; Cokus et al., 2008; Lisch, 2013; Vitte et al., 2014; Eichten et al., 2013). Transposons inserted into new genomic locations may lead to DMRs in the surrounding low-copy sequence though the spreading of DNA methylation from TEs to surrounding sequences (Ahamed et al., 2011; Eichten et al., 2013). To investigate the impact of TE polymorphisms on DNA methylation variation, transposon polymorphisms were identified and compared to the reference Bd21 sequence using the *TEPID* analysis package (Stuart et al., 2016 - Under review; https://github.com/ListerLab/TEPID). Paired-end sequencing data was used from the resequencing efforts of the seven reference *Brachypodium* lines (Gordon et al., 2014) to identify polymorphic TE insertion sites within their respective genomes (see methods). In total, 443 novel TE insertions and 3,576 deletions were identified across the six non-Bd21 genomes (Figure 5A; Sup Table 7). Similar to SNPs and DMRs, Bd1-1 displayed the most insertions and deletions of all tested genotypes, with Bd21-3 and Bd3-1 displaying the fewest. Bd1-1 is the most genetically distant accession of the set compared to Bd21, with an average of 178 bp between SNPs (Gordon et al., 2014). Bd21-3 and Bd3-1 are the most similar with 537 bp and 488 bp between SNPs respectively. Surprisingly, Bd21-3 displays a large number of TE deletions given its close relationship with the Bd21 reference (FIgure 5A). The majority (70%) of TE insertions and many (40%) deletions are present just once in one of the genotypes studied (Sup Fig 15). However, there are examples of transposable elements being inserted multiple times within and/or across different genotypes (Sup Fig 16). A non redundant set of 4,019 transposable element polymorphisms is provided in Supplemental Table 7.

**Figure 5:**
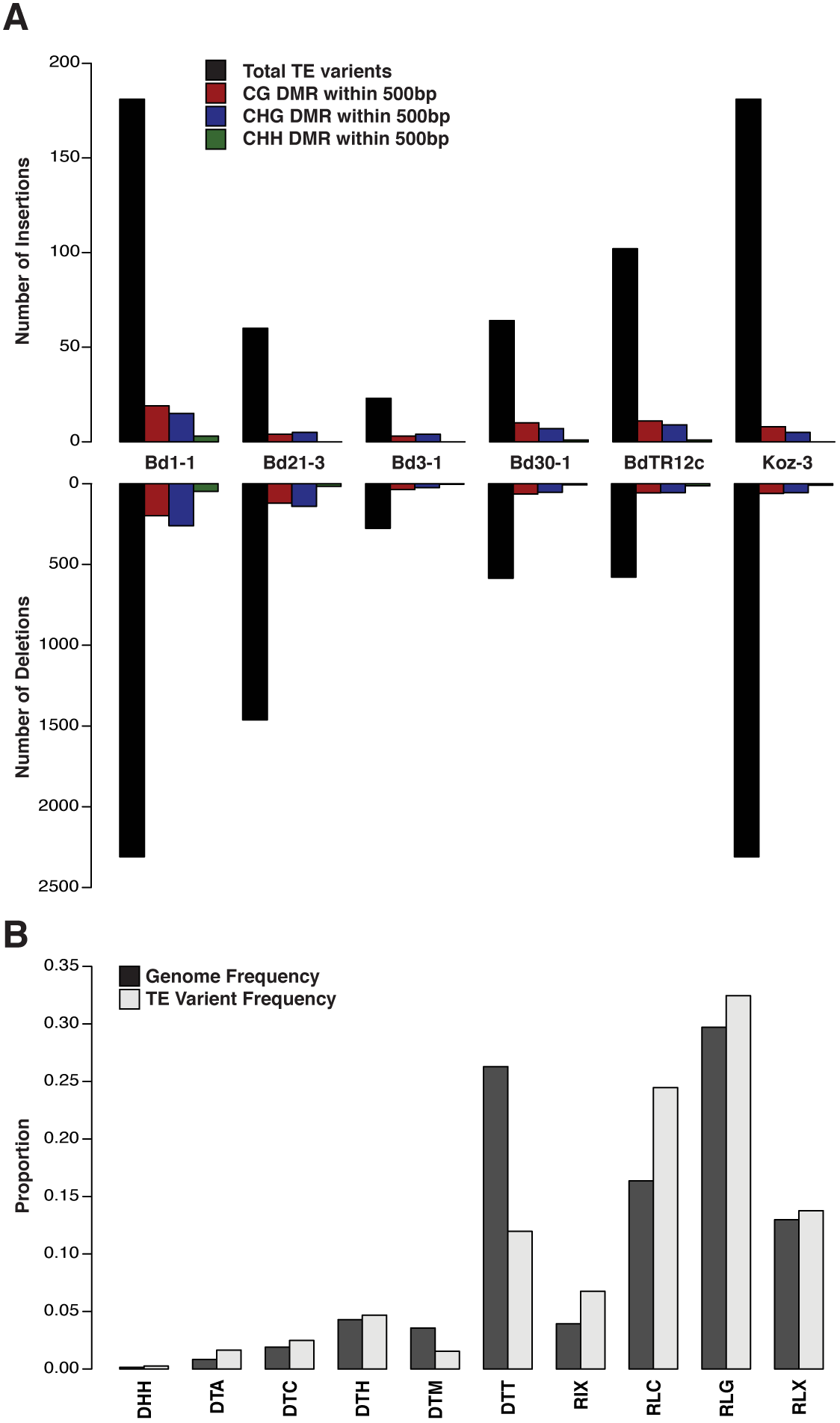
Transposable element polymorphisms across sequenced lines. (A) Barplot indicating total number of transposable element insertions and deletions compared to the Bd21 reference along with the number of DMRs found within 500 bp for each sequence context. (B) Barplot of transposable element families. The genomic frequency of each family (black bars) is compared with the frequency of identified TE insertions and deletion (grey bars).

The transposable element insertions and deletions were also classified based on the TE family which they are derived (Fig 5B). Although the majority of polymorphic TEs are found at a similar distribution to their genomic average, RLC retrotransposons appear to be more polymorphic, and DTT DNA transposons less polymorphic. There were no TE families that did not display at least one insertion or deletion event. Overall hundreds of novel transposon insertions and deletions are identified compared to the reference. Unlike previous studies (Stuart et al., 2016 - Under Review), there appears to be no clear link to DNA methylation variation flanking polymorphic TE sites.

Given the relationship between DNA methylation and transposable elements, CG, CHG, and CHH DMRs identified between each genotype and Bd21 were compared to the sites of TE insertions or deletions to determine the relationship between non-reference TE sites and methylation variation. Possible associated DMRs were filtered to those within 500 bp of a TE polymorphism as likely local associated features. There appears to be minimal enrichment for identified DMRs to be near either transposon insertions or deletions (Figure 5A). This is most prominent in CHH methylation in which almost no TE polymorphic site has a CHH DMR nearby. It would be expected that new TE insertions and deletions would lead to hypermethylation and hypomethylation respectively in the containing genotype. Indeed, a small enrichment for hypermethylation was seen for novel TE insertions as well as hypomethylation for deletions for individual genotypes when compared to the Bd21 reference (Sup Fig 17).

## Discussion

DNA methylation has been shown to associate with a wide variety of annotated features, leading to new insights into the mechanisms of genomic repression of transposable elements, as well as gene regulation (Zilberman et al., 2006; Zemach et al., 2013; Takuno & Gaut, 2013). The DNA methylation profiles of the Bd21 reference genome, as well as six additional resequenced lines (Gordon et al., 2014) of *Brachypodium distachyon*, provide a useful map of DNA methylation across genetically diverse accessions of a model cereal.

Overall, levels of DNA methylation in Brachypodium appear to follow similar patterns to other plant species (Mirouze & Vitte, 2014) in which global methylation level is proportional to repeat content of the genome (Figure 1E). Over half of all CG sites appear methylated across the Bd21 genome (Figure 1A). This is in contrast with CHG and, especially, CHH methylation that are found at much lower levels. These contexts display a similar enrichment for DNA methylation within pericentromeric regions as previously described in plants (Figure 1B; Borowska et al., 2011). Patterns of gene body methylation are similar to those described in the first reported bisulfite sequencing of Bd21 (Figure 1D; Takuno & Gaut, 2013). Curiously, there is limited CHG methylation within gene bodies compared to other monocots such as maize (Sup Fig 7; Eichten et al., 2013). This may be due to the higher repeat content of the maize genome (Figure 1E) and the high proportion of genes containing intronic transposable elements in maize (West et al., 2014).

In comparison, transposable elements of all family types display large amounts of CG and CHG methylation and rarely are found in an unmethylated state. Similar broad patterns have been seen in other plant systems (Zhang et al., 2008). The DTT class of DNA transposable elements displays a somewhat unique pattern of methylation with a proportionally large amount of the genome’s CHH methylation tiles (Figure 1C). This class of Sub1 / Mariner transposable elements is a common feature of the Bd21 genome with over 20,000 annotated elements. DTT elements have been noted for high levels of CHH methylation in other species, as a common target of ‘CHH islands’ defining the boundary between active and repressed chromatin (Gent et al., 2013; Li et al., 2015a). As reported in maize, DTT elements are often found near genes as a common target of CHH methylation (Figure 1D; Sup Fig 18). It is certainly clear that DNA methylation patterns are often unique to specific genomic elements with a wide variety of functions (Diez et al., 2014; Kim & Zilberman, 2014).

Differences in DNA methylation between genotypes may provide insight into genomic regulation and chromatin restructuring. With the DNA methylation profiles of six additional accessions, differentially methylated regions (DMRs) were identified compared to the Bd21 reference (Figure 2A). It should be noted that the vast majority of the genome does not display variable methylation patterns between samples indicating a largely stable chromatin landscape across the species. Indeed, the majority of methylation patterns appear almost identical across annotated features (Sup Fig 9). Even so, variations in DNA methylation levels were observed across accessions, with thousands of DMRs identified across the three methylation contexts studied (Figure 2F). The number of DMRs identified in each accession appears linked to the underlying genetic distance between the lines as determined by re sequencing (Figure 2D-E; Gordon et al., 2014). DMRs in the CG context display ∼30% overlap between accessions, indicating many conserved methylation variants compared to the reference Bd21, which may be the outlier in certain cases (Figure 2B-C). Asymmetric CHH methylation variation appears to often arise from transposable element sequences. As CHH methylation is a *de novo* modification which requires constant targeting, it is possible that certain transposable elements are being actively silenced by the RNA-directed DNA methylation pathway (RdDM; Matzke & Mosher, 2014). In contrast to CHH methylation, CG DMRs are rarely found in transposable elements (Figure 2C). This is similar to previous findings in *Arabidopsis thaliana* (Vaughn et al., 2007; Schmitz et al., 2013). As a repressive mark targeting fully-silenced transposons (Kim & Zilberman, 2014), it is unlikely that CG DMRs would arise from regions of the genome fully marked for heterochromatic silencing. Given the order of magnitude difference in symmetric (CG and CHG) compared to asymmetric (CHH) methylation, it is clear that unique pathways (such as RdDM) are major drivers of the methylation state of these DMRs. Therefore, the ability to analyze these DMRs separately is of clear importance.

Biological replication of DNA methylation profiles has largely been lacking to date. By investigating DNA methylation across replicate samples of Bd21, Bd1-1, and Bd3-1 we found high levels of correlation for CG methylation between replicates (Figure 3A). This correlation was slightly weaker for CHG and substantially less for CHH (FIgure 3B-C). Previous reports in replicated Bd21 bisulfite libraries have indicated a high level of variability between replicates (Roessler et al., 2015). Our data largely supports this for CHH methylation as correlation levels are quite low when compared to other types of genomic assays across replicates (e.g. RNA seq biological replicates r2 ∼ 0.99; Makarevitch et al., 2015). Symmetric DNA methylation sites are captured often by both single, and multiple replicate data (Figure 3E), however there are many DSS-based DMRs that appear to be missed by the tile approach (Figure 3F). Depending on the questions to answer, one may prefer to have a larger number of DMRs as possible candidate variants (combined with a larger chance of false positives). This is in contrast to the conservative approach of multiple filters required for the tile-based DMRs. The two DMR approaches also highlight the variability of CHH methylation across samples and the methods used to identify it. There was almost no overlap between CHH DMRs between methods (Figure 3E). This lack of conservation between methods may indicate a fundamental limitation in the DMR identification methods as used. However, it also may be tied to the minimal correlation between biological replicates for this sequence context (r2 ∼0.35) in that there is not likely to be much overlap. As CHH methylation is continually established *de novo*, it may not show the same level of biological stability when compared to the symmetric methylation contexts which have maintenance methyltransferases to maintain fidelity over DNA replication (Law & Jacobsen, 2010).

A major question regarding the study of DNA methylation is the likelihood that observed patterns are either dependent on genetic state, or act independently as a separate epigenetic layer of regulatory information (Richards, 2006; Eichten et al., 2014). By looking at high quality SNPs across the reference lines sequenced (Gordon et al., 2014), DNA methylation variation was correlated with increased levels of genetic variation (Figure 4). For all three contexts, correlation values from 0.42-0.56 were observed indicating that a proportion of all DMRs identified across these diverse lines are likely tied to the genetic variation found nearby. Although this correlation is observed, it does not eliminate the possibility of unlinked DNA methylation variation that acts independent of genetic state. The results show that the relationship between genetics and DNA methylation is clearly complex (Eichten et al., 2014) with some, but not all, DNA methylation variation associated with genetic states.

A possible genetic source of DNA methylation variation may be transposable elements as they are known to be the major target of DNA methylation that acts to suppress their activity (Kim & Zilberman, 2014; Mirouze & Vitte, 2014). It is possible that variation in transposable element content could create novel targets for DNA methylation and lead to differential methylation between samples (Eichten et al., 2012; Mirouze & Vitte, 2014; Stuart et al., 2016-Under-Review). An analysis of paired-end sequencing data of the six resequenced lines identified hundreds of novel transposable element insertions and deletions compared to the Bd21 reference (Figure 5A). Evidence in maize and Arabidopsis has suggested that many DMRs within the genome may be tied to the presence or absence of certain transposable elements (Hollister & Gaut, 2009; Hollister et al., 2011; Ahmed et al., 2011; Eichten et al., 2012; Eichten et al., 2013). Surprisingly, there is minimal evidence for DNA methylation variation surrounding novel transposable element insertions or deletions (Figure 5A). It is possible that if some transposon polymorphisms are recent events, they may not be targeted for heterochromatin silencing. It is also possible that these insertions or deletions may be occurring in regions that are already highly heterochromatic, leading to minimal changes in overall DNA methylation patterns in the surrounding regions and limiting sequencing coverage of repetitive regions.

The landscape of DNA methylation and transposable element polymorphisms within diverse accessions of *Brachypodium distachyon* indicate chromatin variation that is often tied to underlying genetic variation. However, there was no clear evidence to tie novel transposable element polymorphisms to nearby DNA methylation variation. Although examples of transposable element presence-absence variation influencing methylation state has been reported (Hollister & Gaut, 2009; Hollister et al., 2011; Ahmed et al., 2011; Eichten et al., 2012; Eichten et al., 2013; Stuart et al., 2016), other reports indicate minimal association between transposable element variation and methylation (Li et al., 2015b) which may indicate species specific relationships. Given the relationship between DNA methylation state and genetic background, it would be of interest to investigate natural populations with less genetic variation between them. Preventing large population structure may assist further studies to identify the activity of these methylation variants and novel insertions in relationship to possible functional consequences such as transcriptional regulation.

## Methods

### Tissue collection

Seeds were germinated in moist petri dishes for one week at 10°C and transferred to soil. Plants were grown under 12hr light conditions at 18-21°C in controlled growth rooms. Three week old mature leaf tissue was harvested from each of the seven inbred *B. distachyon* lines.

### Brachypodium Bisulfite Sequencing

gDNA was extracted from harvested tissue using Qiagen DNAeasy Plant kit (Limberg, Netherlands) and quantified using the Qubit HsDNA (Life Technologies, Carlsbad, CA). 50ng of purified DNA was bisulfite converted using the Zymo DNA- Gold bisulfite conversion kit (Zymo Research, Irvine, CA). Whole genome bisulfite sequencing (WGBS) libraries were constructed using the EpiGnome Post-Bisulfite Library Kit (Epicentre, Madison WI) per manufactureres instructions (EPILIT405 rev. C). Libraries were quantified on the Perkin Elmer GXII and Agilent BioAnalyzer (Santa Clara, CA) to confirm library quality. Libraries were subsequently pooled and sequenced (Paired-end, 100bp) across a HiSeq 2000 (Illumina, CA) lane with a 10% PhiX control DNA spiked in for cluster control.

### Read alignment and methylation calling

The resulting reads (Sup Table 1) were trimmed to remove adapter contamination and poor quality reads using TrimGalore (http://www.bioinformatics.babraham.ac.uk/projects/trim_galore/). Trimmed reads were then aligned to SNP corrected versions of the Bd21 reference genome (Gordon et al., 2014) using Bismark (v0.12.5, Krueger & Andrews, 2011). As Epignome PBAT libraries appear to create a large number of chimeric reads, alignment had to be performed in three stages to maximize mapping efficiency. First, a traditional directional paired-end alignment through Bismark was performed (flags: -n 2 -l 20 --un). The unmapped ‘Read 1’ reads were then processed through a directional single-end alignment in Bismark (flags: -n 2 -l 20). The unmapped ‘Read 2’ reads were also processed through directional single-end alignment with the additional ‘--pbat’ flag allowing mapping to the complementary strands. The three resulting alignments were run through the bismark methylation extractor (flags: --comprehensive --report --buffer_size 8G) in their expected paired-end or single-end modes. Paired-end methylation extraction included the ‘--no_overlap’ flag to prevent counting the same cytosine if covered by both the forward and reverse read. Output was then merged by sequence context (CG, CHG, CHH) and run through bismark2bedGraph (flags: --CX). 100bp tiled windows providing proportion methylated, number of methylated reads, and number of unmethylated reads across the genome were also created for downstream analysis

### DMR identification

The number of DMRs identified in this study are similar to other studies in plants. However, the various methods and filtering criteria that are used largely inhibit direct comparisons between lists. The described DMRs in this study are largely filtered to be a conservative estimate of variable methylation sites by requiring strict read count, size, and differential methylation levels and biological reproducibility.

Differentially methylated regions between Brachypodium samples was performed using a novel pipeline based on 100bp tiled windows across the genome. In brief, for each pairwise sample comparison, all windows were called differentially methylated if the absolute difference in proportion methylation met a given threshold (CG: 80%; CHG 50%; CHH 20%). Windows were then filtered to require at least 10x coverage across the window to be valid. All adjacent windows were collapsed into a single DMR region. All results were compared and the largest region was kept for any overlapping DMRs between pairwise comparisons. All DMRs used for analysis were mapped to annotated genes (Bdistachyon_192) and genomic repeats (Brachy_TEs_V2.2; IBI 2010) using Bedtools (Quinlan & Hall, 2010).

### Replicate methylation data

Tissue from five Bd21, five Bd1-1, and four Bd3-1 plants was harvested, DNA extracted, and libraries prepared as described above. 100bp Single-end sequencing on the HiSeq2000 (Illumina, CA) was performed with a 10% PhiX control DNA spiked in for cluster control. DMRs were called using DSS without smoothing and q-value below 0.01. All other parameters were kept as default. The comparisons are done pairwise with respect to all three lines and sequence context (CG, CHG, CHH).

### Identification of Transposon polymorphisms

Brachypodium paired-end reads sequenced by Gordon et al. (2014) were mapped to the SNP-corrected reference for each accession using bowtie2 (Langmead et al. 2012) and yaha (Faust and Hall, 2012), using the mapping script included in the TEPID package with insert sizes estimated from Gordon et al. (2014). TE polymorphisms were then identified by running ‘tepid-discover’ using the Brachypodium TE annotation from Gordon et al. (2014), included in the TEPID package, and the ‘--mask’ option set to mask all scaffold chromosomes. TE insertion calls were then refined and accessions genotyped for each variant using ‘tepid-refine’.

### SNP density calculation across genome

The overlapping set of SNPs between SOAP and MAQ as identified in Gordon et al (2014) were used for this analysis. SNPs were subset to those found between Bd21 and each of the six accessions independently and binned by genomic location into 1Mb (CG, CHG) or 5Mb (CHH) windows with no overlap. DMRs for each accession and sequence context were binned in a similar fashion using Bedtools (Quinlan and Hall, 2010).

### Methylation levels of other plant species

Whole genome bisulfite sequencing sequence data of other plant species (Sup Table 8) were collected from the sequence read archive of associated publications (Lister et al., 2008; Chodavarapu et al., 2012; Eichten et al., 2013). Reads were processed with TrimGalore and Bismark with the same parameters as *Brachypodium* samples. Reads were mapped as either paired-end or single-end given availability. Sup Table 5 lists all samples, alignment metrics, and reference genomes used for non Brachypodium analyses.

## Data Access

All sequencing reads have been deposited under SRX993729 in the NCBI Short read archive (SRA). Scripts and pipelines for sequencing alignment and DMR calling are available at www.github.com/borevitzlab/wgbs_analysis. Transposable element polymorphisms calling source code can be found at https://github.com/ListerLab/TEPID.

## Acknowledgements

The authors would like to acknowledge the work of the Australian National University Biomedical Research Facility for sequencing support as well as John Vogel regarding *Brachypodium distachyon* annotations and resequencing efforts. SRE was supported by an Australian Research Council Discovery Early Career Research Award and Human Frontiers Science Program LTF fellowship. TS was supported by the Jean Rogerson Postgraduate Scholarship.

### Author Contributions

SRE and JOB designed experiments. RL and JOB supervised experiments. SRE performed sequencing. SRE, TS, and AS performed analyses. SRE wrote manuscript

### Author ORCID IDs

0000-0003-2268-395X (SRE)

0000-0002-3044-0897 (TS)

---

**Supplemental Figure 1:**
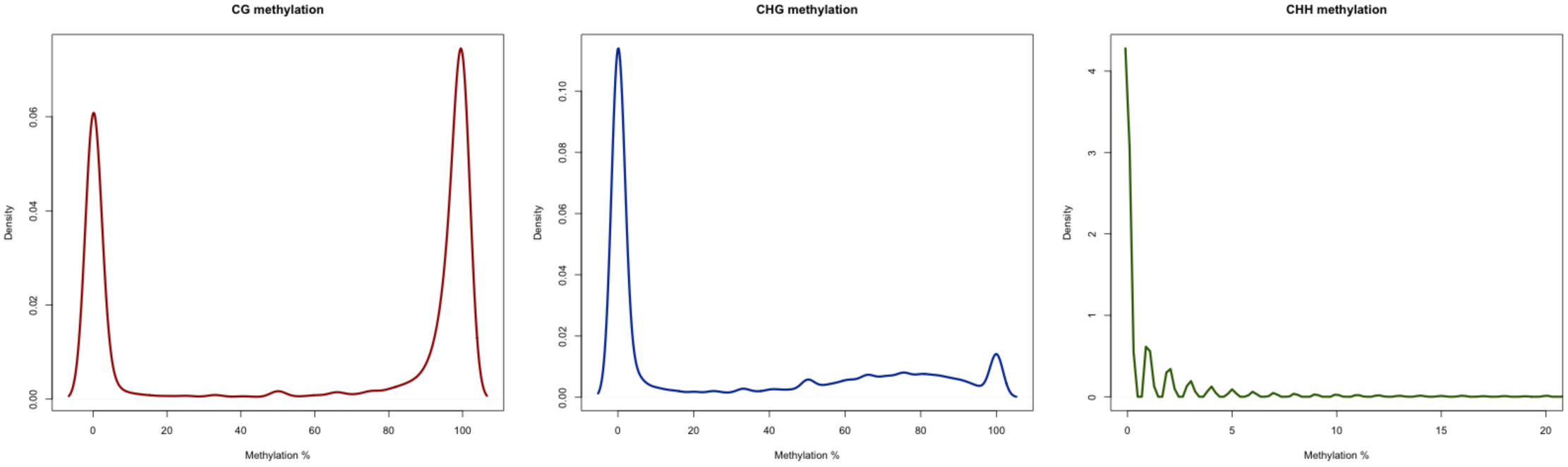
Density distributions describing CG, CHG, and CHH methylation Brachypodium.

**Supplemental Figure 2:**
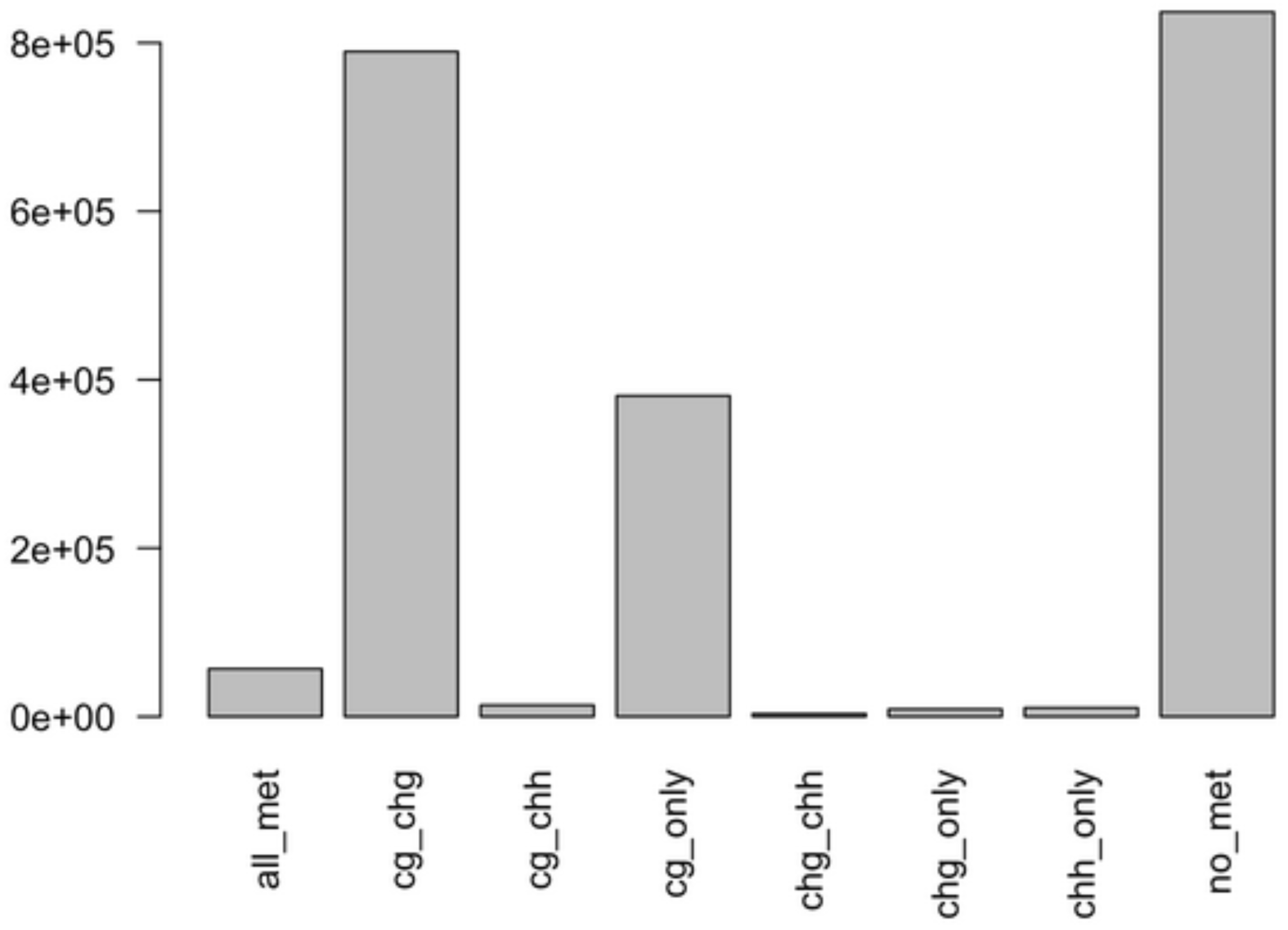
Number of genomic tiles falling into eight possible DNA methylation classes in Bd21

**Supplemental Figure 3:**
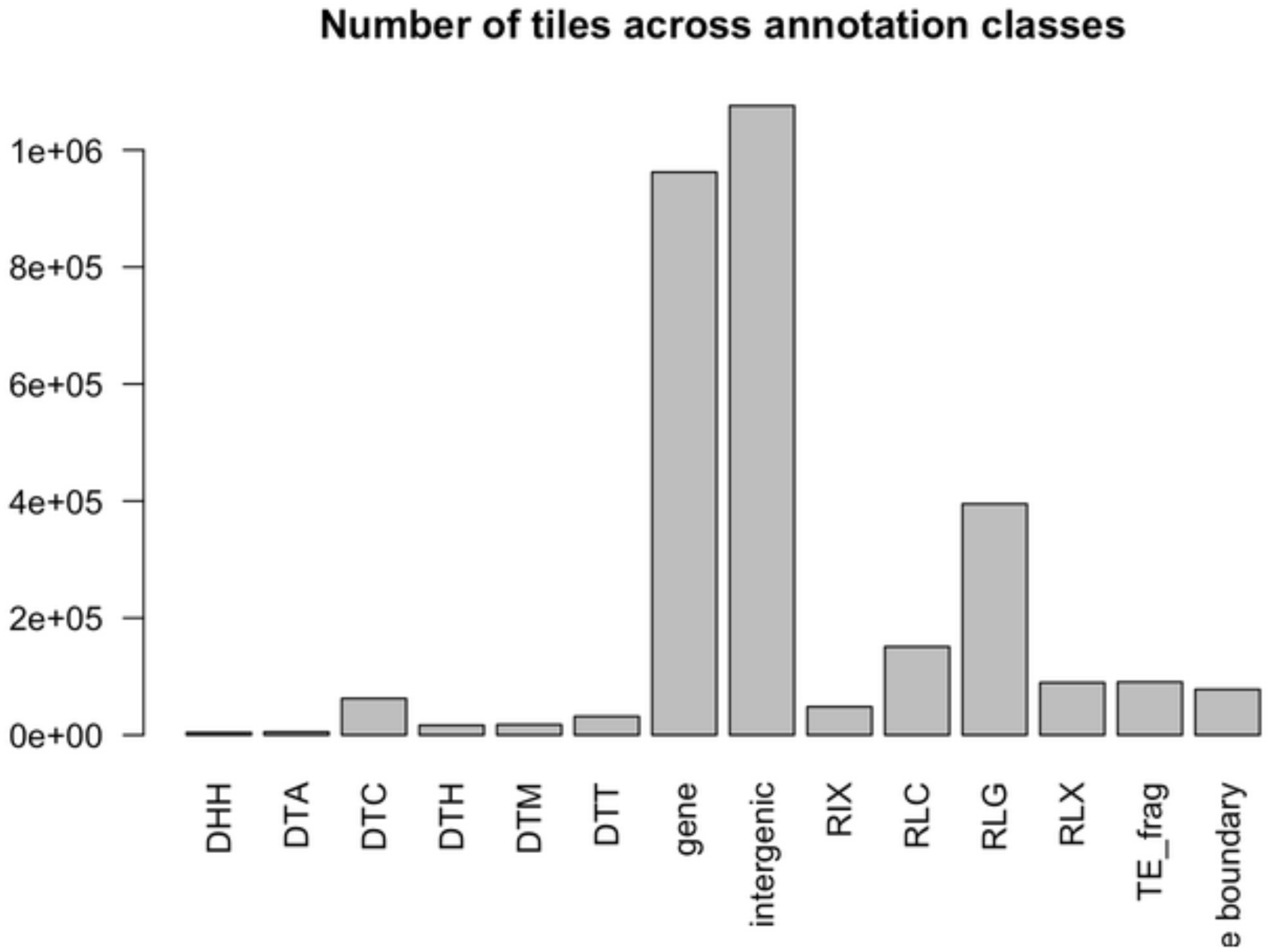
Number of genomic tiles intersecting with annotation features

**Supplemental Figure 4:**
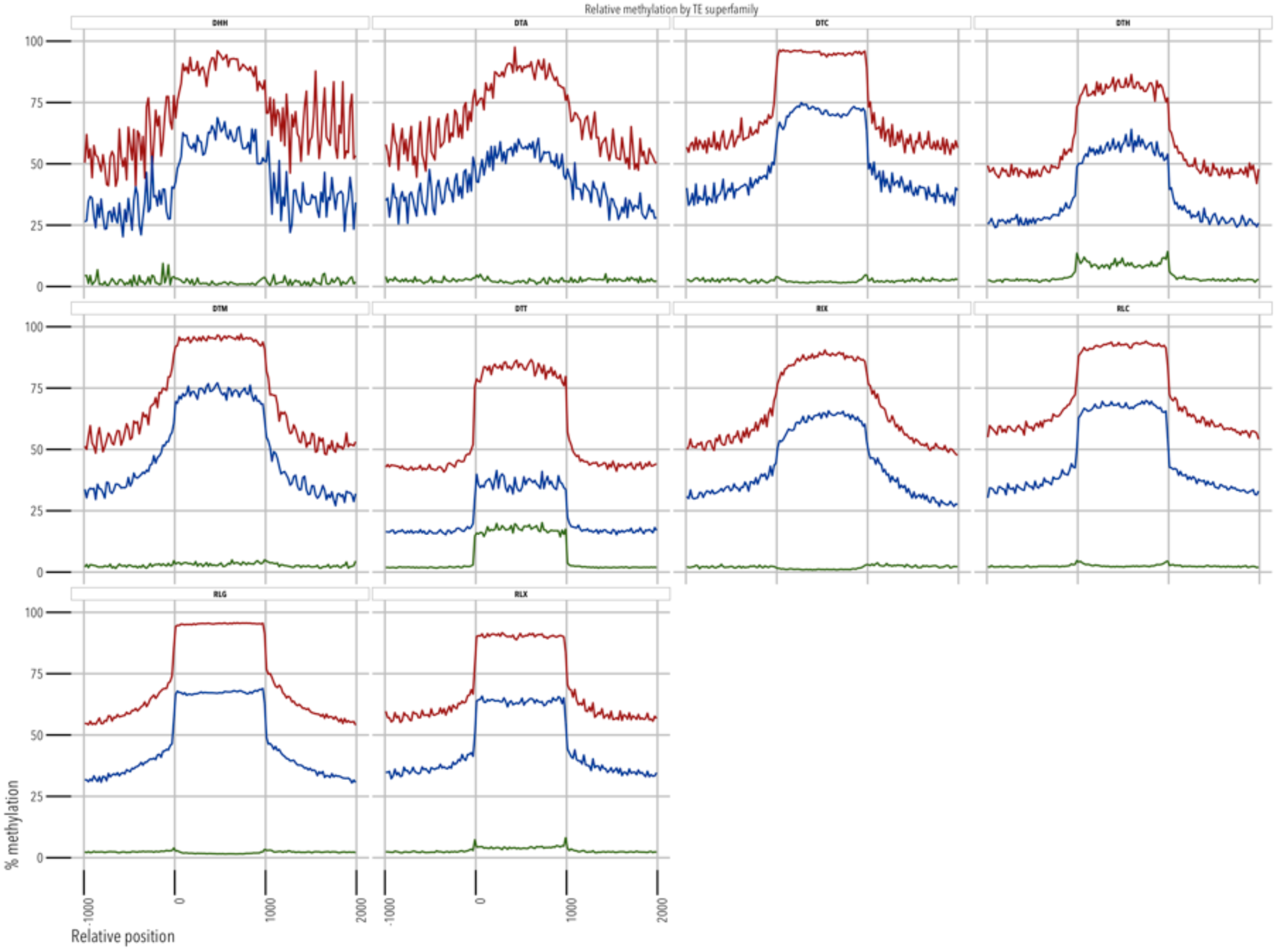
Relative distance plots of average methylation for ten *B. distachyon* annotated transposon superfamilies. Methylation is shown in CG (red), CHG (blue), and CHH (green) contexts.

**Supplemental Figure 5.**
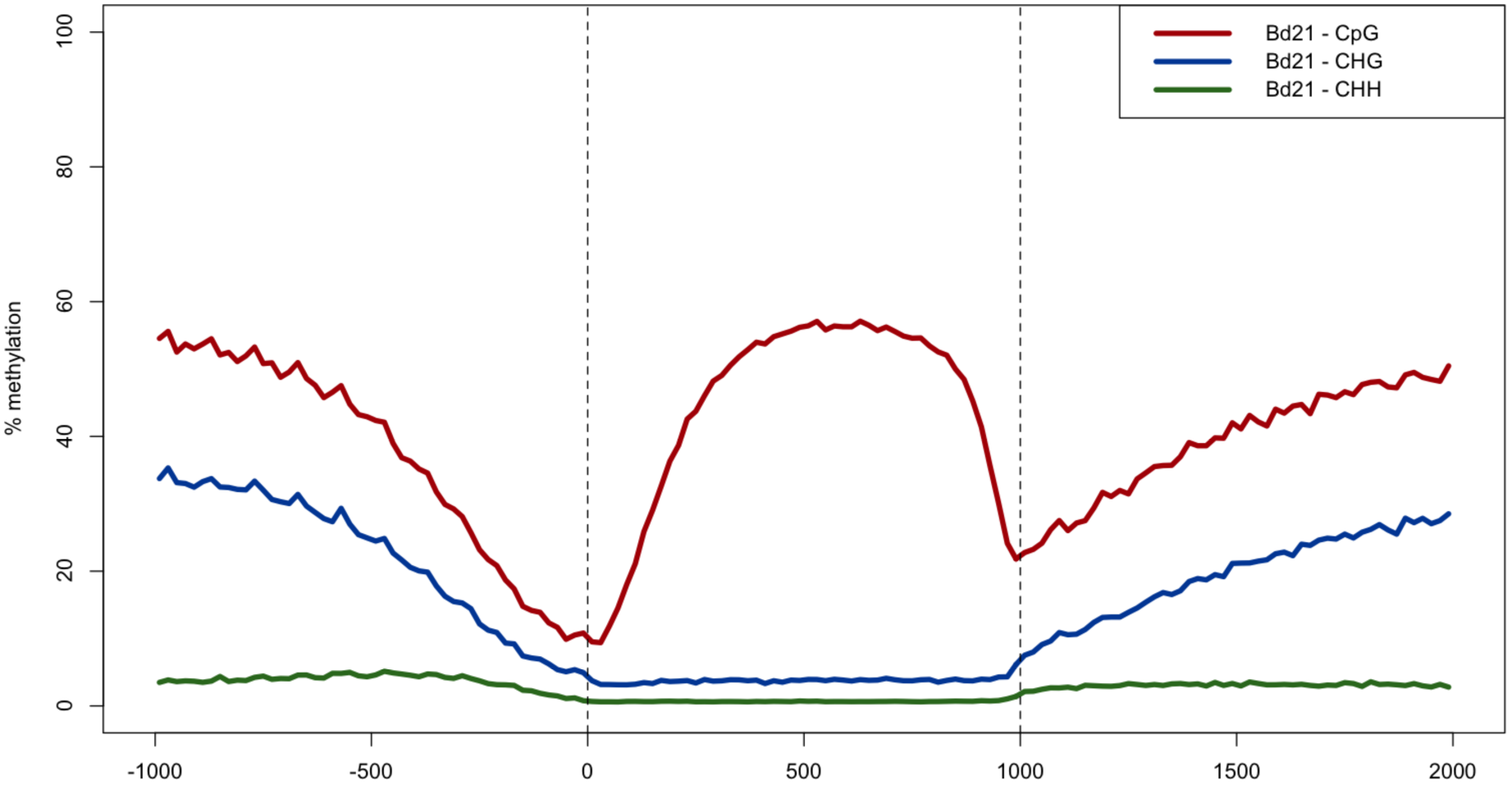
Relative methylation of annotated Bd21 genes with intronic gene sequences removed. CG gene-body methylation is maintained without intronic sequence methylation levels.

**Supplemental Figure 6:**
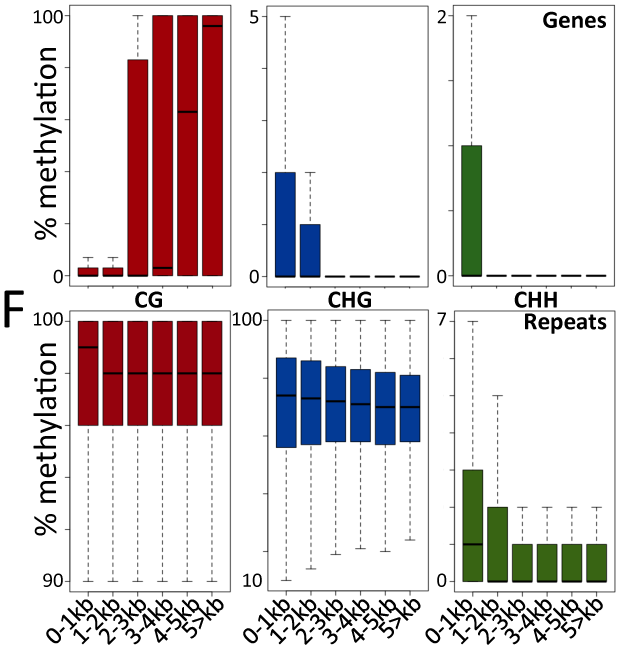
DNA methylation for genes (top) and TEs (bottom) divided by element size

**Supplemental Figure 7:**
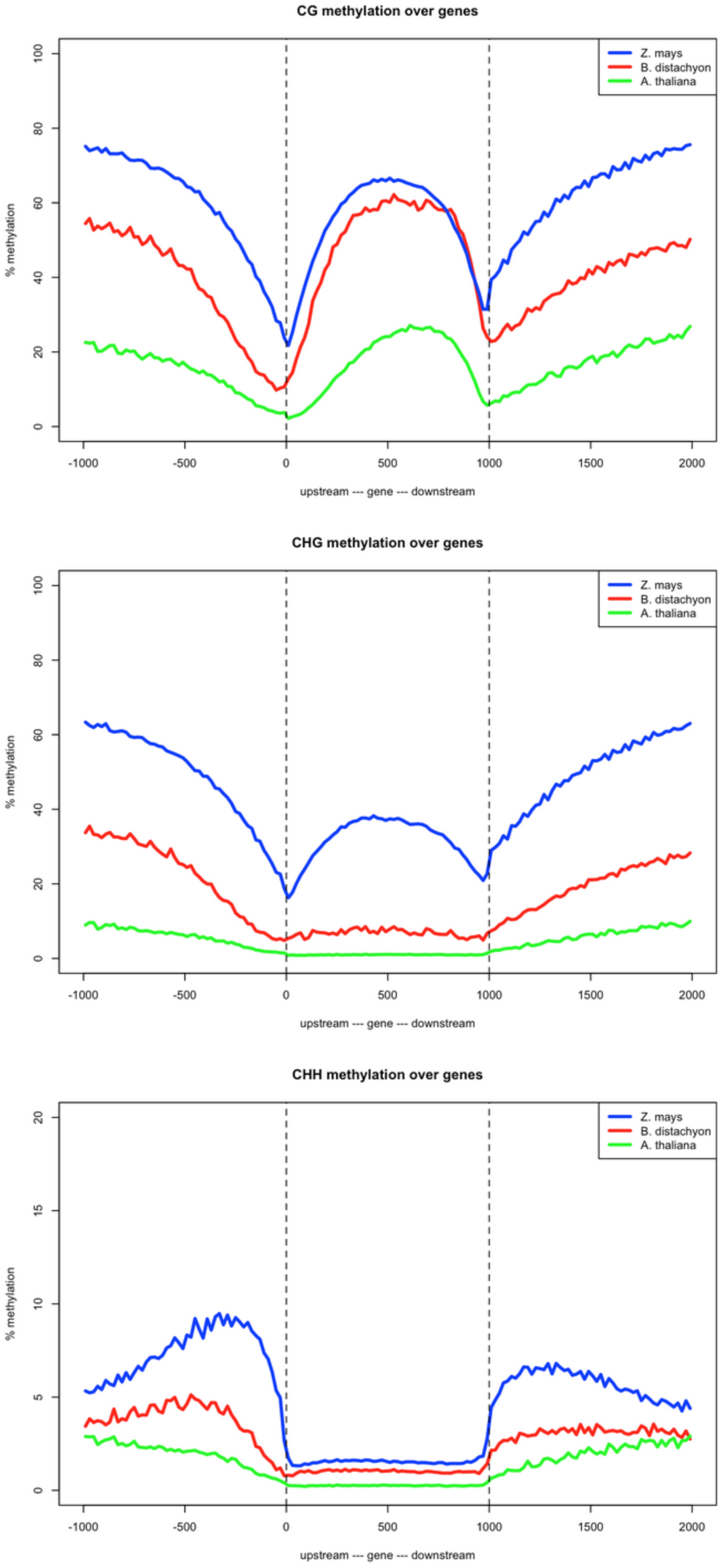
Methylation comparisons of B73 (Z. mays), Col-0 (A. thaliana), and Bd21 (B. distachyon) over gene models. Plots are colored by species and split by methylation sequence context. Note scale for CHH methylation plot

**Supplemental Figure 8:**
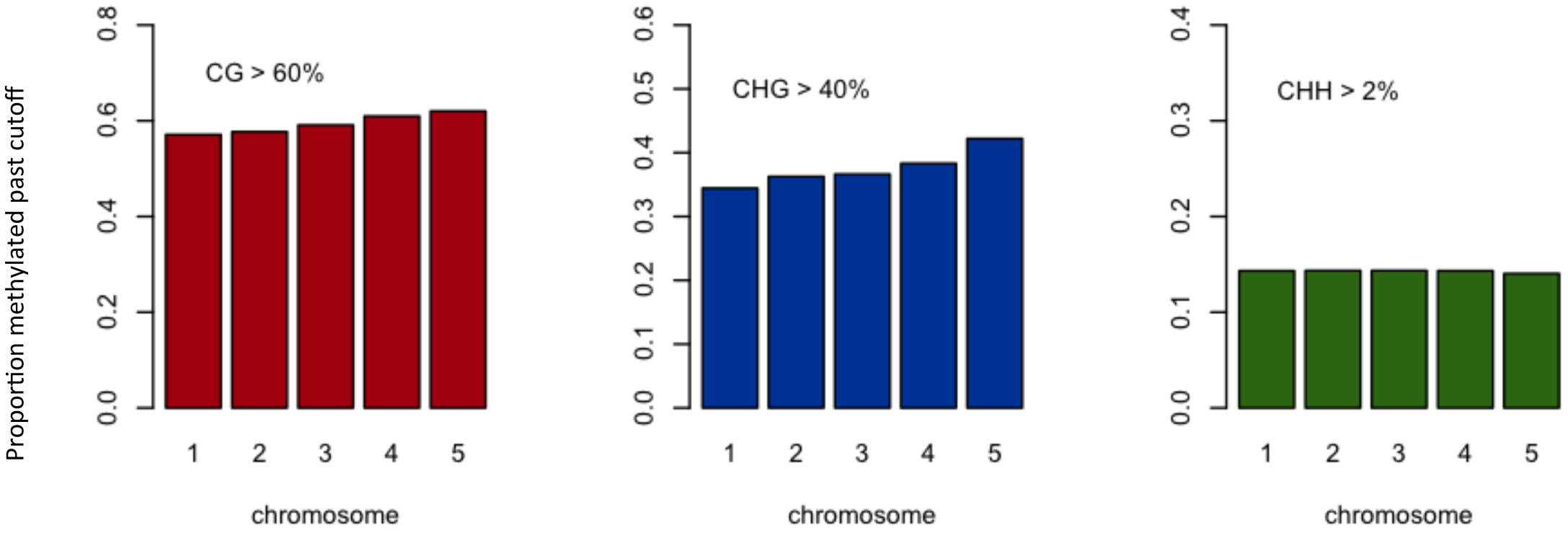
Proportion of methylated tiles across Bd21 chromosomes. Percentages defining methylated are provided

**Supplemental Figure 9.**
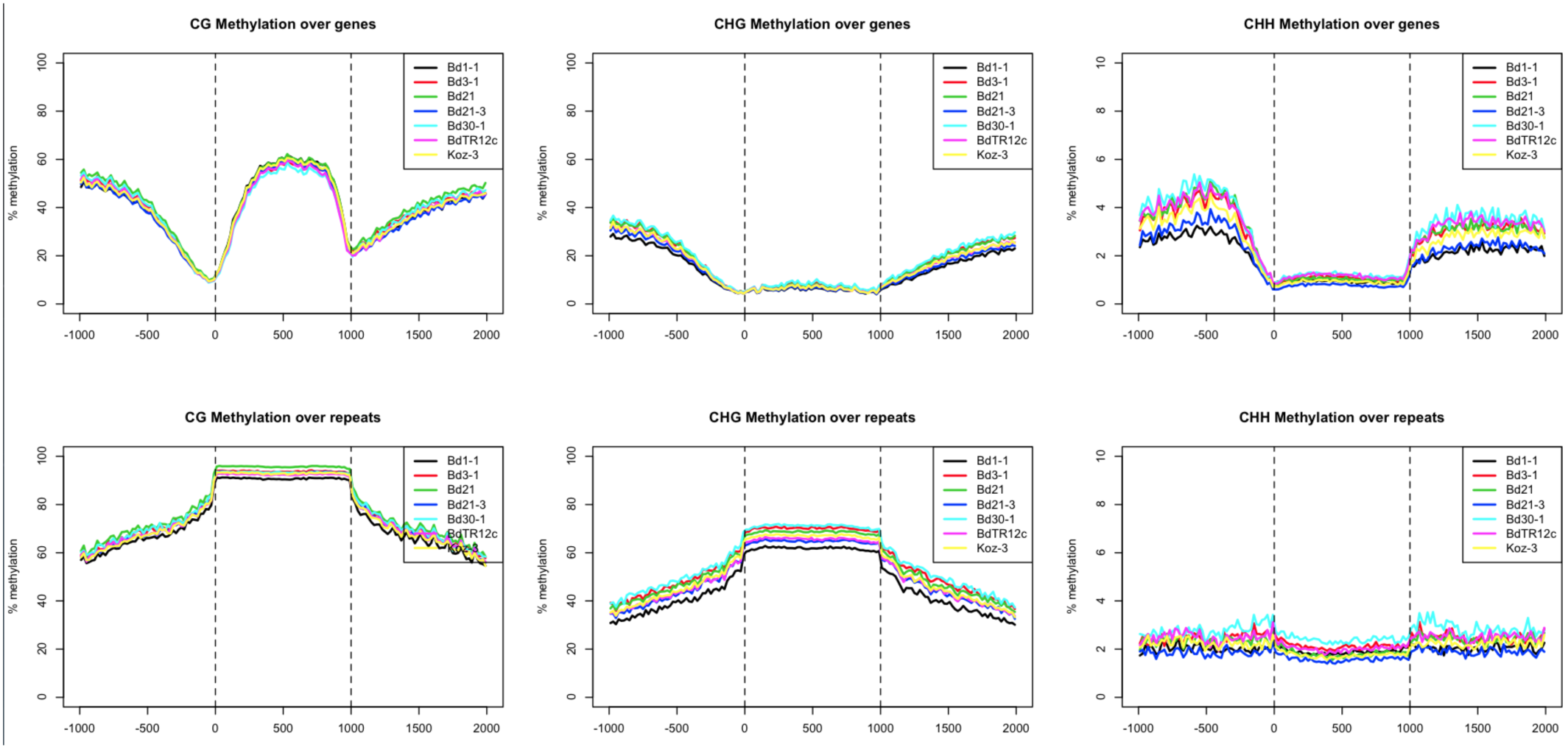
Relative methylation across genes and repeats for all seven inbreds. Plots have been split by methylation context.

**Supplemental Figure 10.**
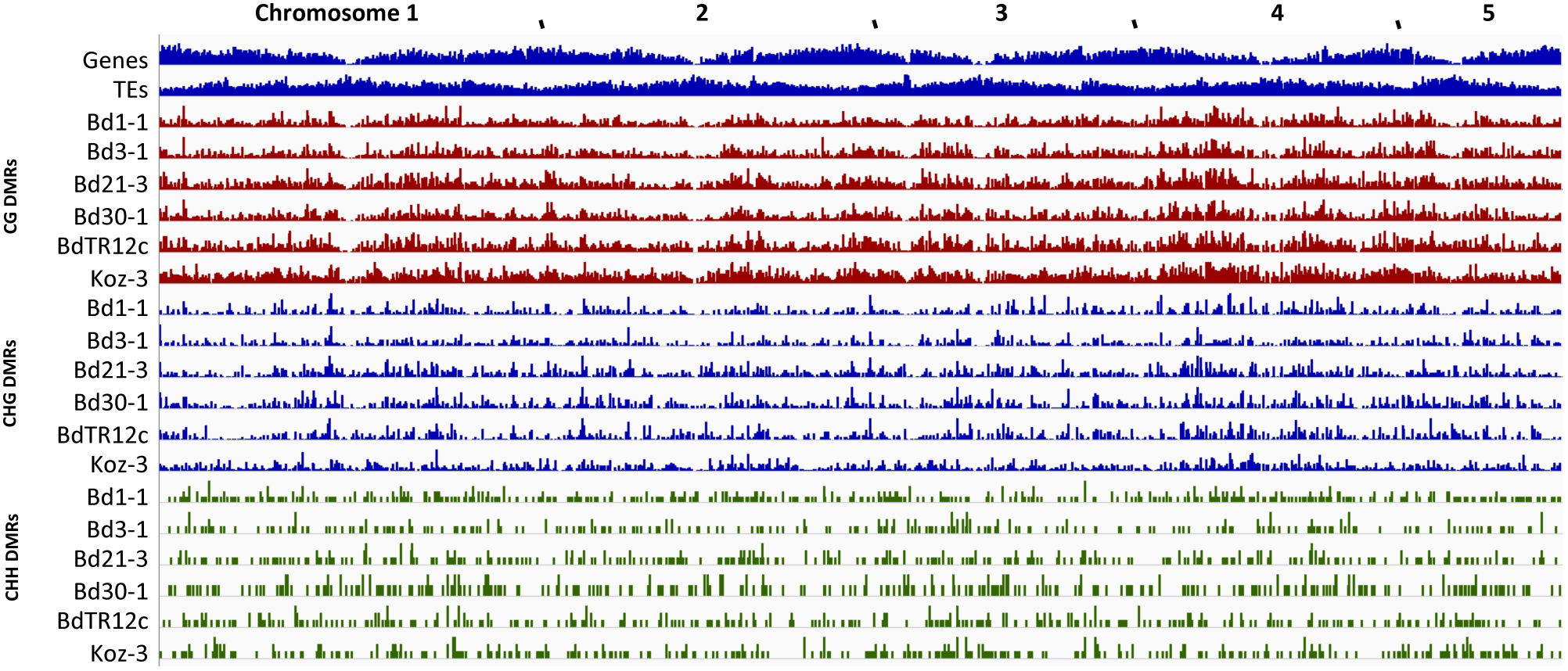
Position of DMRs across all lines and chromosomes

**Supplimental Figure 11.**
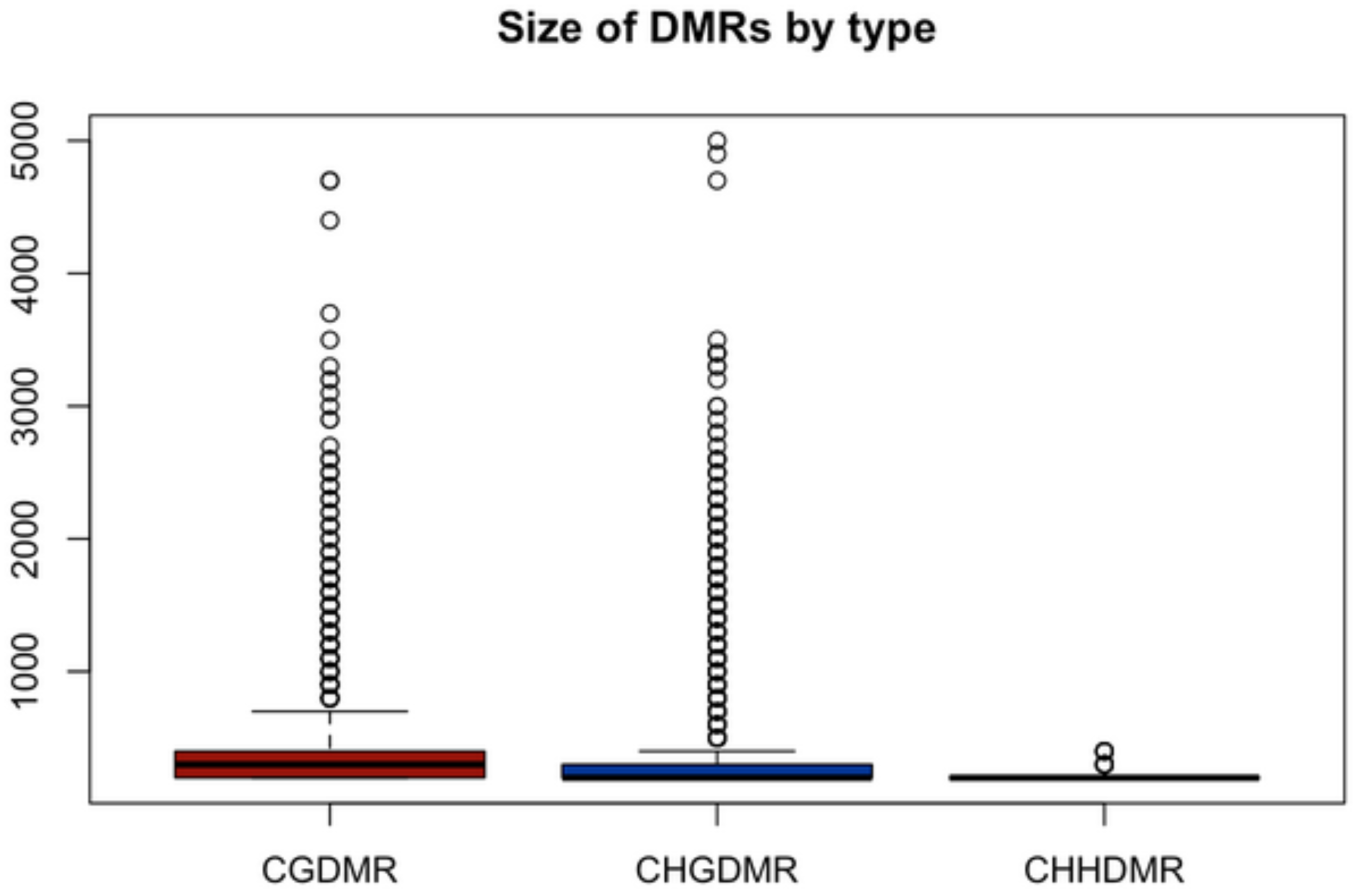
Boxplot of DMR sizes per sequence context

**Supplimental Figure 12.**
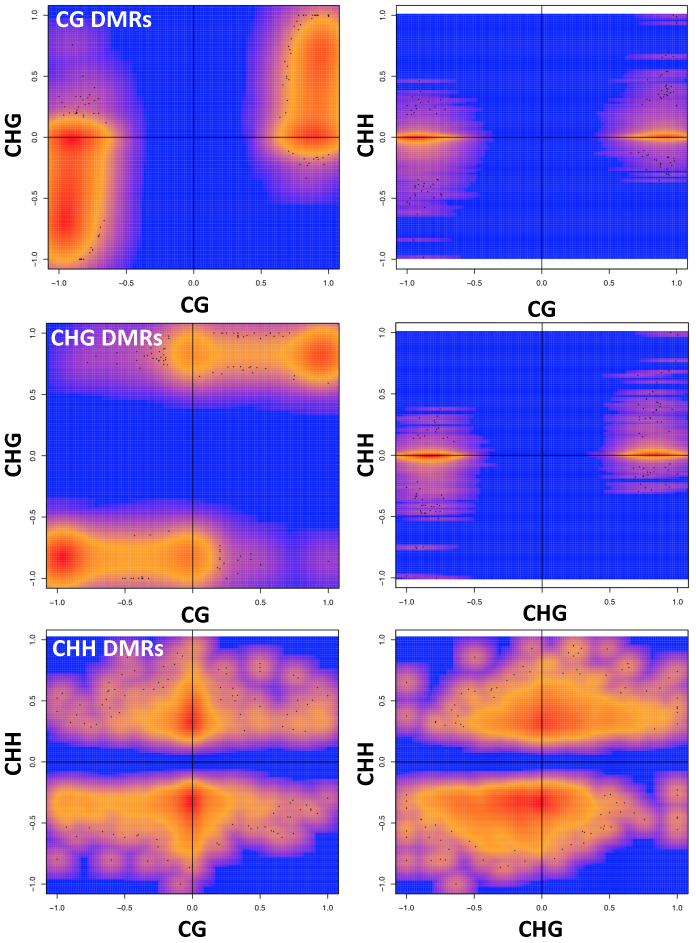
Scatter plots of methylation states across CG, CHG, and CHH DMR regions. Red indicates increased point cloud density.

**Supplemental Figure 13.**
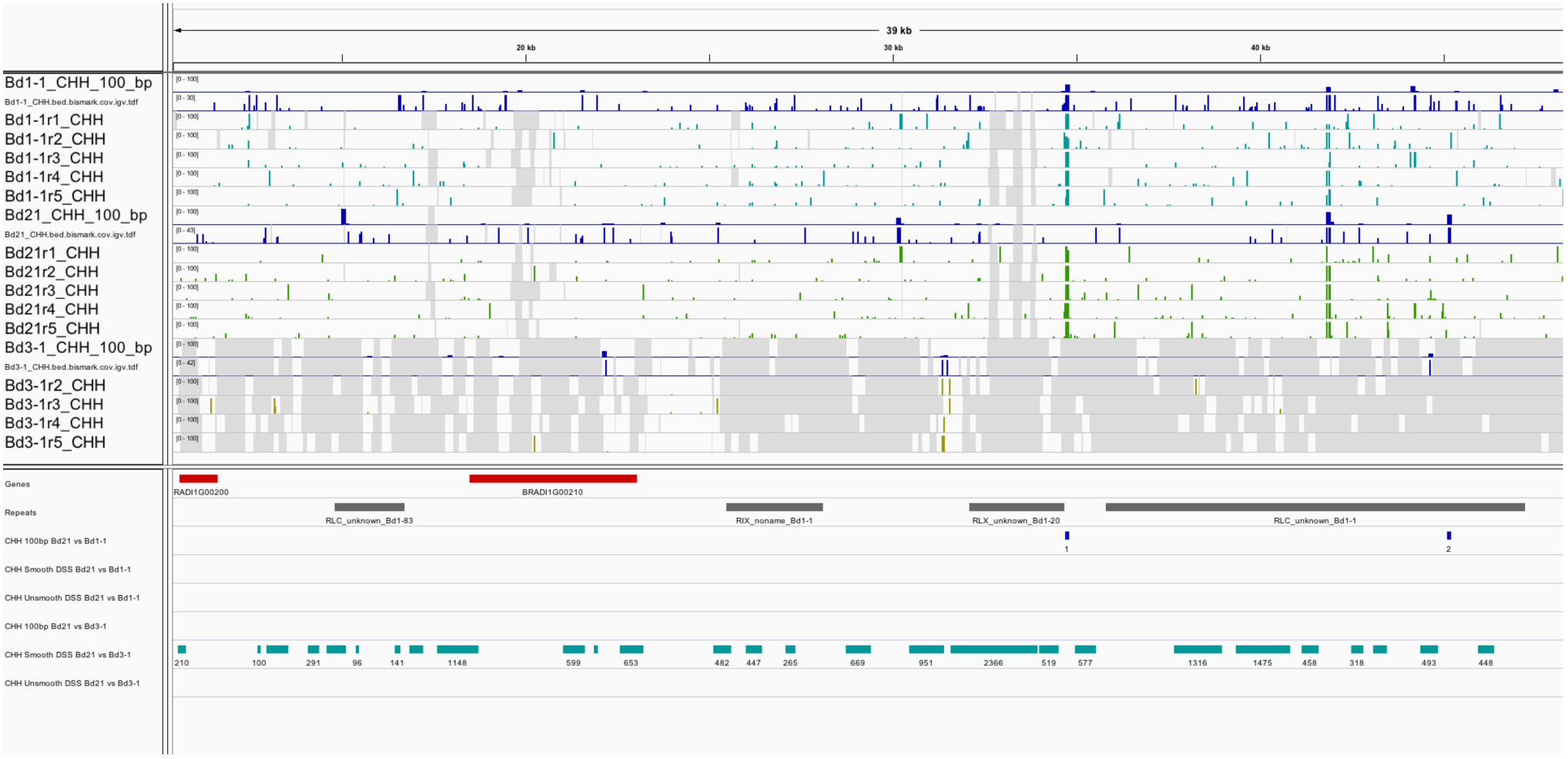
Genomic view of Bd1:10,391-49,958 highlighting smooth DSS DMRs being called across a largely absent region of Bd3-1. Aqua bars indicate DMR calls.

**Supplemental Figure 14:**
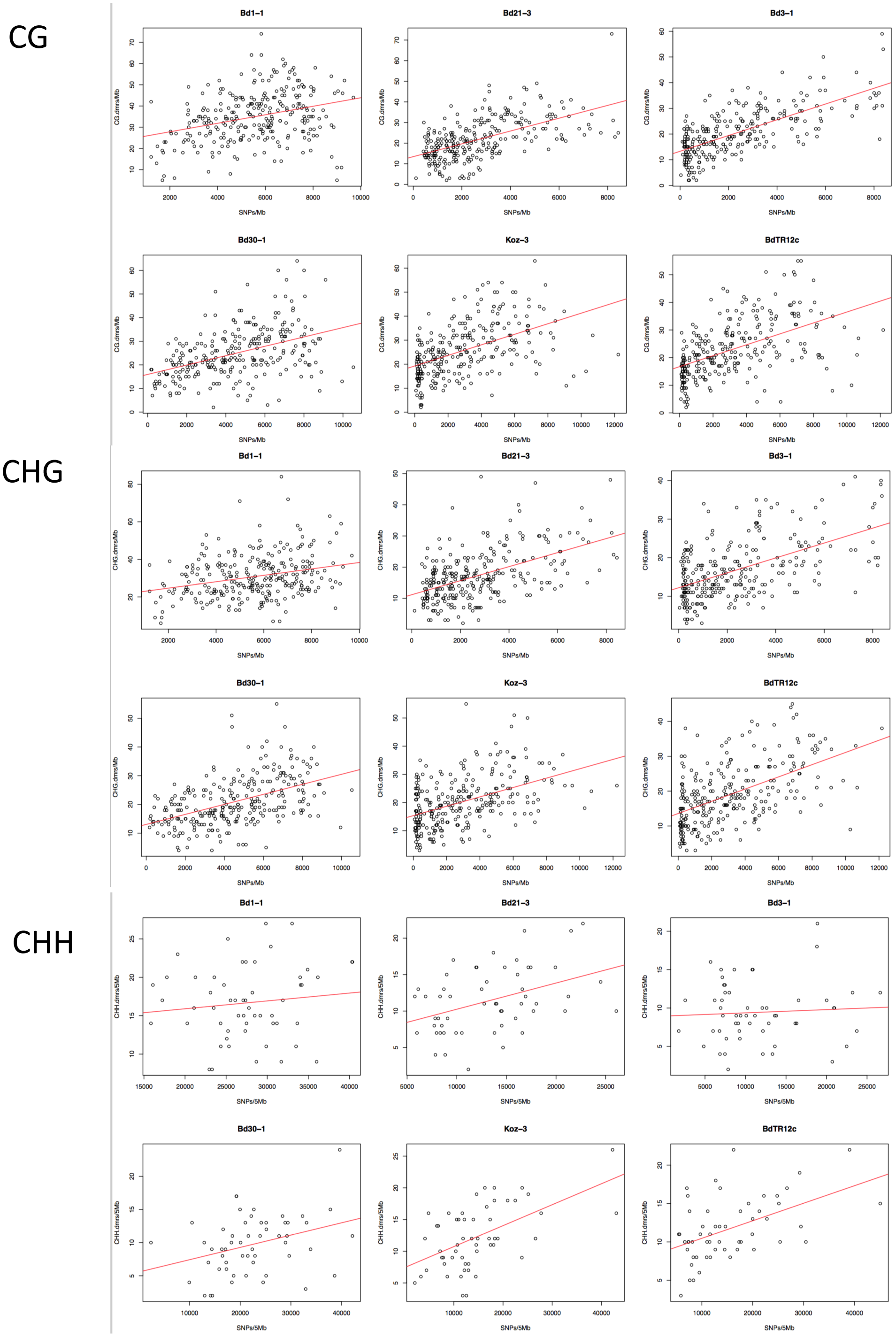
Correlation plots for all DMR types for all individual samples

**Supplemental Figure 15.**
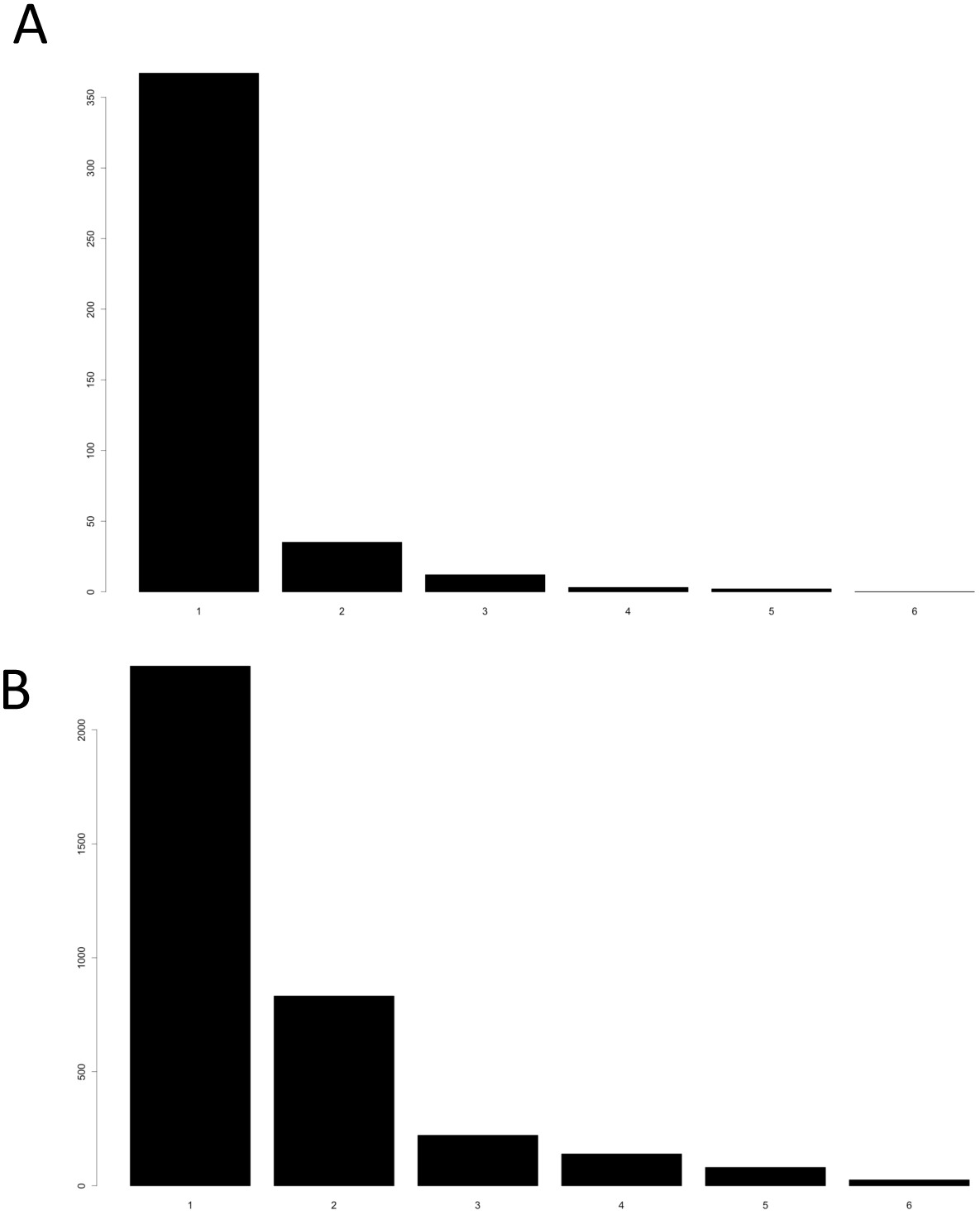
Barplots indicating the number of genotypes individual transposable elements are inserted (A) or deleted (B) across samples.

**Supplemental Figure 16.**
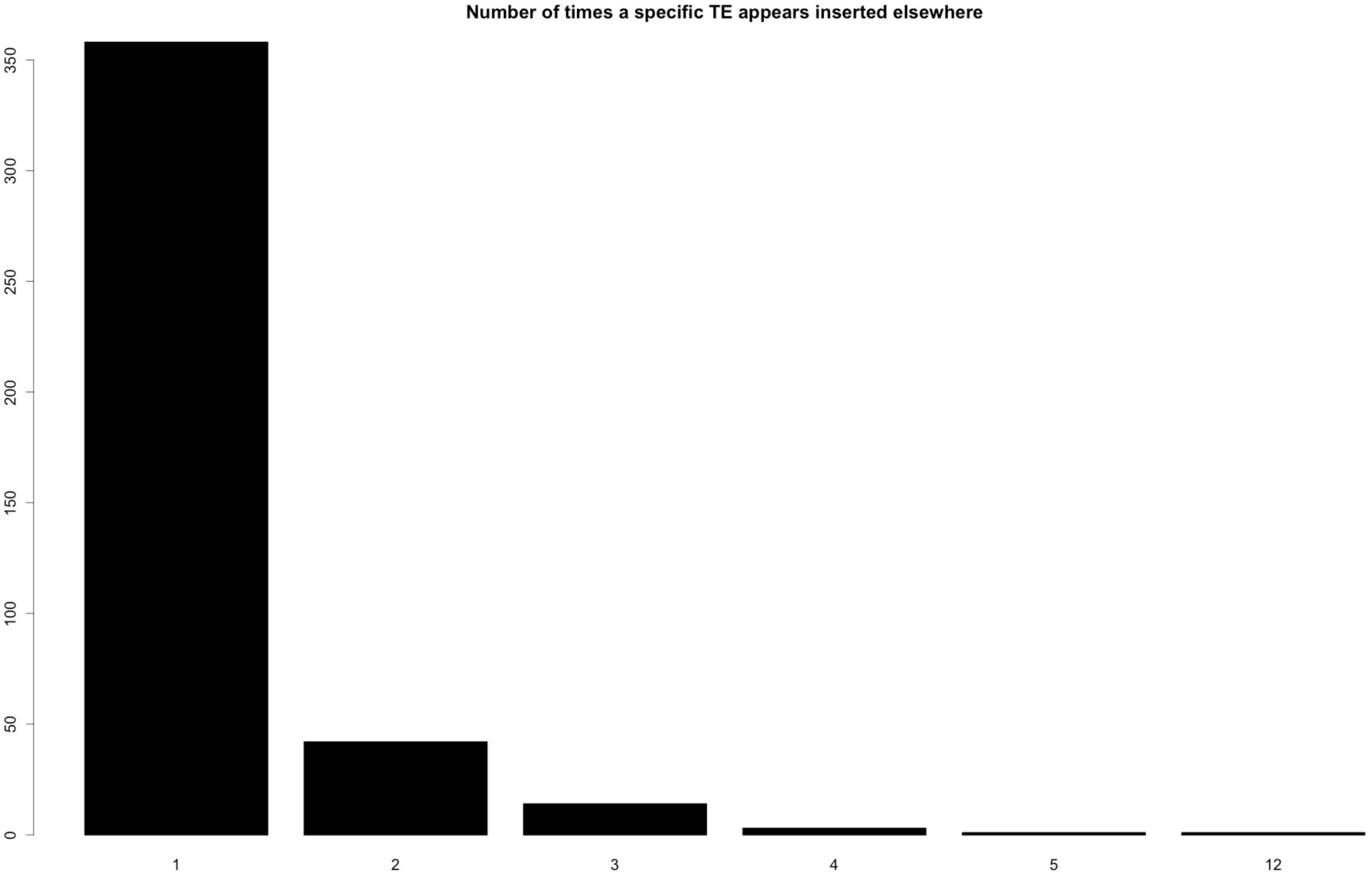
TEs inserted into multiple genotypes and/or multiple times in a single sample

**Supplemental Figure 17.**
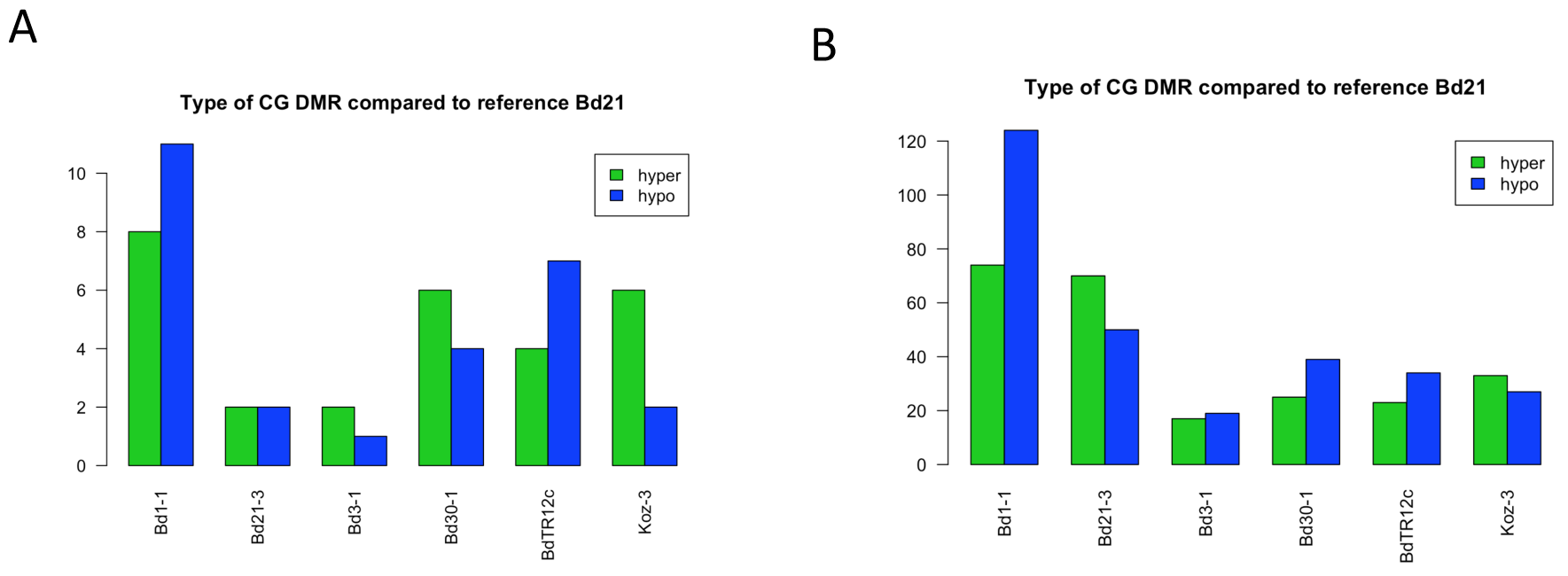
Barplots indicating the DMR state (hyper or hypoethylated) compared to the Bd21 reference methylation state for DMRs within 500bp of (A) transposable element insertions and (B) deletions

**Supplemental Figure 18.**
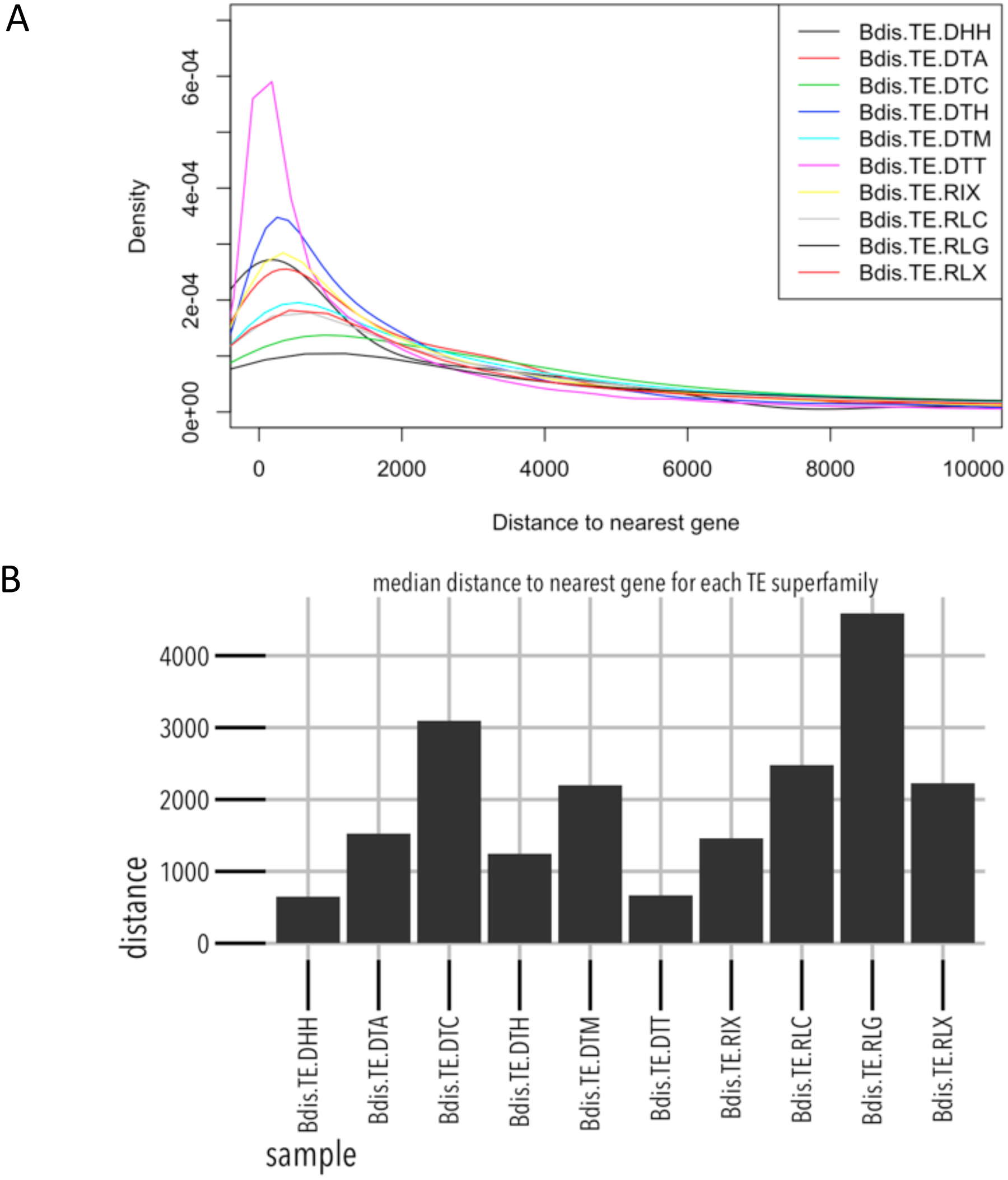
Density distribution (A) and median distance (B) of each annotated transposon superfamily to its nearest gene.

**Supplemental Table 1:** Bisulfite sequencing summary statistics

**Supplemental Table 2:** List of CG/CHG Differentially Methylated Regions

**Supplemental Table 3:** List of CHH Differentially Methylated Regions

**Supplemental Table 4:** Bisulfite sequencing summary statistics for biological replicate data

**Supplemental Table 5:** List of Smooth DSS DMRs

**Supplemental Table 6:** List of Unsmooth DSS DMRs

**Supplemental Table 7:** Transposable element polymorphisms in Brachypodium distachyon

**Supplemental Table 8:** Bisulfite sequencing summary statistics for other plant species

